# Modelling coordinated nocturnal rhythms in ungulates

**DOI:** 10.1101/2024.09.13.612804

**Authors:** Johann Ukrow, Jennifer Gübert, Max Hahn-Klimroth, Paul Wilhelm Dierkes, Gaby Schneider

**Author notes:** Robert-Mayer-Str. 10, 60325 Frankfurt, Goethe-University Frankfurt, Germany (Gaby Schneider).

## Abstract

Studying animal behavior is an important aspect of ethology and behavioral biology and a prerequisite to improving animal management in zoos. As the nocturnal behavior of many ungulate species is, in contrast to the behavior during daylight, poorly studied, a better understanding of nocturnal behavior is necessary to improve animal welfare.

We analyse the nocturnal behavior of ungulates recorded in a large number of German and Dutch zoos. These animals show a switching between standing and lying phases, which can be associated with a certain degree of regularity. Interestingly, this regularity is not always captured in the simple length distributions of behavioral phases but shows in the autocorrelation and in the coordination of standing and lying across animals. Particularly, this phenomenon often occurs in younger animals.

We provide an explanation to this phenomenon by proposing a stochastic model that can describe these processes. For individual behavior, regular standing cycles are assumed to be potentially interrupted by short lying phases, such that a regular background rhythm appears in the autocorrelation but not necessarily in the raw length distribution of the activity phases. For coordinated behavior, crosscorrelation functions allow to analyse the degree to which pairs of animals that are sharing the same stable box show a synchronization of their standing-lying rhythms.

In the data set, our analyses suggest that indeed, baseline regularity does not seem to be reduced in younger animals. Instead, younger animals showed increased probabilities for interruption of standing phases by short lying phases. In addition, the coordination of the standing-lying rhythm between animals in the same box ranged up to 100% and decreased with the distance between boxes. We also found systematic delays between the standing activity of young and adult animals.

**Author summary:** In this paper, we investigate rhythms observed in the nocturnal behavior of a large number of ungulates housed in zoos. Motivated by the observation that younger individuals, in particular, exhibit irregular standing-lying cycles, we propose and analyze a mathematical model to describe the basic rhythms of the observed animals and quantify the regularity of their standing-lying behavior.

Our model enables us to measure both the regularity and synchronization of behavior among individuals. The results suggest that even individuals initially perceived as highly irregular actually follow a strict base rhythm during the night, but single standing phases may be interrupted. Moreover, we observe that animals stalled together typically synchronize their behavior, particularly in the case of a dam and her calf.

## 1 Introduction

The ungulate orders artiodactyla and perissodactyla comprise over 250 species combined [1], many of which are iconic animals of the African landscape. Consequently, numerous widely kept zoo animals belong to these orders [2]. Therefore, understanding these animals’ behaviors is crucial, both for scientific advancement and for improving animal management and welfare. Indeed, knowledge of an animal’s behavioral patterns provides insights into its requirements and needs, making it a prerequisite for enhancing husbandry practices, ultimately improving animal welfare [3–6].

Decades of ethological research have focused on the behavior and daily rhythms of various ungulates. Particularly daily rhythms have been studied, among others, for European Bisons (*Bos bonasus*) [7], Hippotraginae [8], Plains Zebras (*Equus quagga*) [9], Water Deer (*Hydropotes inermis*) [10], and Gerenuks (*Litocranius walleri*) [11]. However, most ungulates of the orders perissodactyla and artiodactyla are diurnal or crepuscular [12] which means that behavioral patterns vary strongly between daytime and nighttime [12–14]. Hence, comprehensive understanding of ungulate behavior requires nocturnal studies as well.

While some species like Giraffes (*Giraffa camelopardalis*) [15–17] and African Elephants (*Loxodonta africana*) [13] have been extensively studied for their nocturnal behavior, there is limited literature on many other ungulates in this regard.

One particular recent line of research presented an ethological description of the nocturnal behavior of a variety of ungulates kept in European zoos [18–21]. Studying the basic behavioral patterns of zoo animals offers two main advantages over studies conducted in the wild. First, zoos provide easier and continuous access to the animals, allowing to collect more data than would be possible otherwise [22]. Second, observing zoo animals allows to directly draw conclusions about improving housing and animal welfare, as deviations from typical behavior can indicate stress [16].

### 1.1 Aims and plan of the paper

The current study builds upon and extends the findings of [21]. This study investigated, among other things, *lying cycles*, defined as the period from when an animal lies down to the next time it lies down, analogous to the well-known sleep cycles in human biology [23]. In [21], the typical length of lying cycles and their distribution over the nights were presented, primarily to detect differences based on taxonomy, age, or sex. It occurred that most adult animals seemed to show a high degree of rhythmicity in the sense that lying cycle length followed approximately a normal distribution. However, some animals, especially young individuals, deviated strongly from this typical pattern and exhibited a bimodal distribution of lying cycle length, with an unexpectedly high number of short lying cycles. This motivated the following questions:

First, can one describe the *regularity* in an individual’s behavior, and do young individuals really exhibit less regularity in their behavior compared to adults? Second, if several individuals show regular standing-lying behavior, do they also synchronize their rhythmic behavioral patterns, and if so, can this synchronization be quantified? Additionally, if an animal with regular behavior and another with seemingly irregular behavior are close to each other, to what degree do their behavioral patterns synchronize?

To approach these questions, we propose a mathematical model to describe this rhythmicity. For a more direct biological interpretation of the behavioral synchronization, we switch from lying cycles to the equivalent standing cycles (SCs), defined as the periods between two successive events of standing up (Figure 1 A). The SCs exhibit phenomena analogous to the LCs, with approximately normal distributions in regular cases (Figure 1 B), bimodal distributions with a remarkably high fraction of short SCs in seemingly irregular cases (Figure 1 C), and a certain synchronization of standing phases among animals sharing a stable box (Figure 1 D).

**Fig 1.**
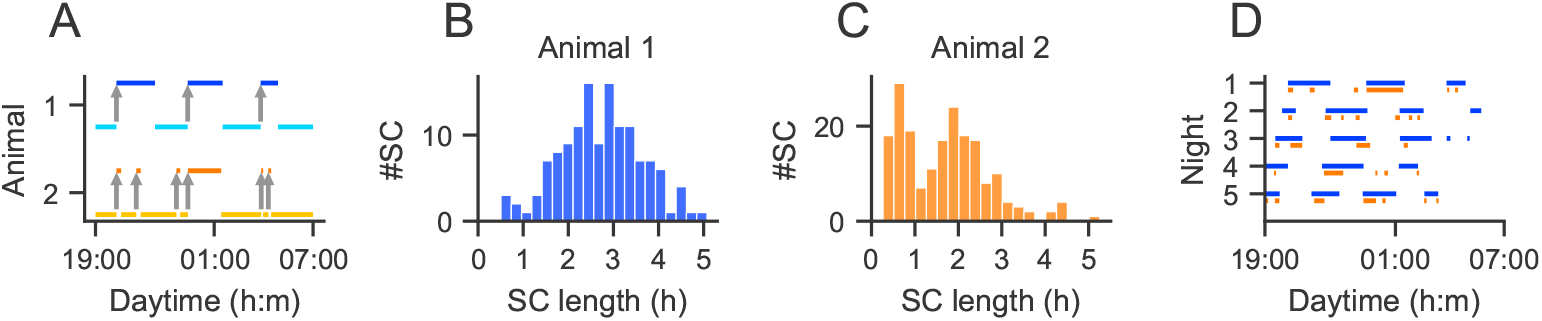
Standing cycles (SC) and their distributions. A SC is defined as the period between two successive events of standing up (grey arrows). **A**. Regular standing-lying behavior of one female adult *Connochaetes taurinus* (animal 1) during one night (blue) and irregular standing-lying behavior of one female young *C. taurinus* (animal 2) during the same night (orange). **B, C**. Unimodal distribution of all SCs of animal 1 (B) and bimodal distribution of animal 2 (C) across all nights. **D**. Standing behavior of animal 1 (blue) and animal 2 (orange) during five nights in the same stable box.

First, we propose a simple model called the *Beta Normal model* (BN, Section 2.2), which is used to describe the typical regular SCs using a few simple assumptions. The BN can be applied to individuals with regular SCs but not to those with seemingly irregular, bimodal SC distributions. In the latter case, the BN fails to capture the high proportion of very short SCs, and cannot reproduce certain regularities observed in the autocorrelation function (ACF). Therefore, we propose a simple extension of the BN, maintaining its simple and regular structure while introducing a stochastic notion of *interruptions* of standing cycles as follows. We allow animals to interrupt the standing phase and to *return* to the same standing phase after a while. This model will be called *Interrupted Beta Normal model* (IBN, Section 2.3). Interestingly, the IBN naturally produces the distributions of SC lengths observed in [21] while keeping the underlying regular structure.

Second, to analyze the degree of synchronization between pairs of animals’ behavioral patterns, we extend the IBN to pairs of processes (Section 2.4), assuming that the times of standing up occur synchronously in both animals. Based on this model, we use the *crosscorrelation function* (CCF) to analyze the degree of synchronization between pairs of animals’ behavioral patterns, as a function of the distance between their stables in the zoo.

Our analyses of a large data set of 9156 single nights from 192 individuals across 19 ungulate species suggest that even behavioral patterns of individuals, which appear *seemingly* irregular at first glance, actually follow a regular base rhythm. Furthermore, the results suggest that animals that share a box may synchronize their standing-lying behavior and that the degree of synchronization decreases with the distance of the animals in the zoo building. In addition, the model also allows for the investigation of systematic temporal shifts in standing up and lying down between adult and young animals.

## 2 Models and methods

### 2.1 The data set

The present article is based on the data set reported in [21]. In short, behavioral data were collected by video recordings with night vision cameras. Classification of behavior was performed using the deep learning-based software package BOVIDS (Behavioral Observations by Videos and Images using Deep Learning Software, [24]). Among the postprocessing steps, standing and lying sequences shorter than 5 min were discarded, and OUT periods were minimized. Animals were classified into juvenile, subadult and adult, where the boundaries were defined as the species’ time of weaning and sexual maturity, respectively. For details on the recording and postprocessing see [21].

### 2.2 The Beta Normal model (BN) for regular individual nights

To describe regular SC distributions as shown in Figure 1 A (upper part) and B, we assume in the BN that SC lengths are independent and normally distributed (conditioned on being positive, i.e., truncated at zero), and that the fraction of lying during each standing cycle is independent and beta distributed (Figure 2 A). The formal definition is provided below in Definition 2.1. Section 3.2 will discuss the model assumptions and demonstrate high agreement with the properties of the present data set. Note that the truncated normal distribution is used because the degree of truncation is usually small (see Figure 1 B) and other distributions restricted to positive values would not simplify the formal notation in later sections.

**Fig 2.**
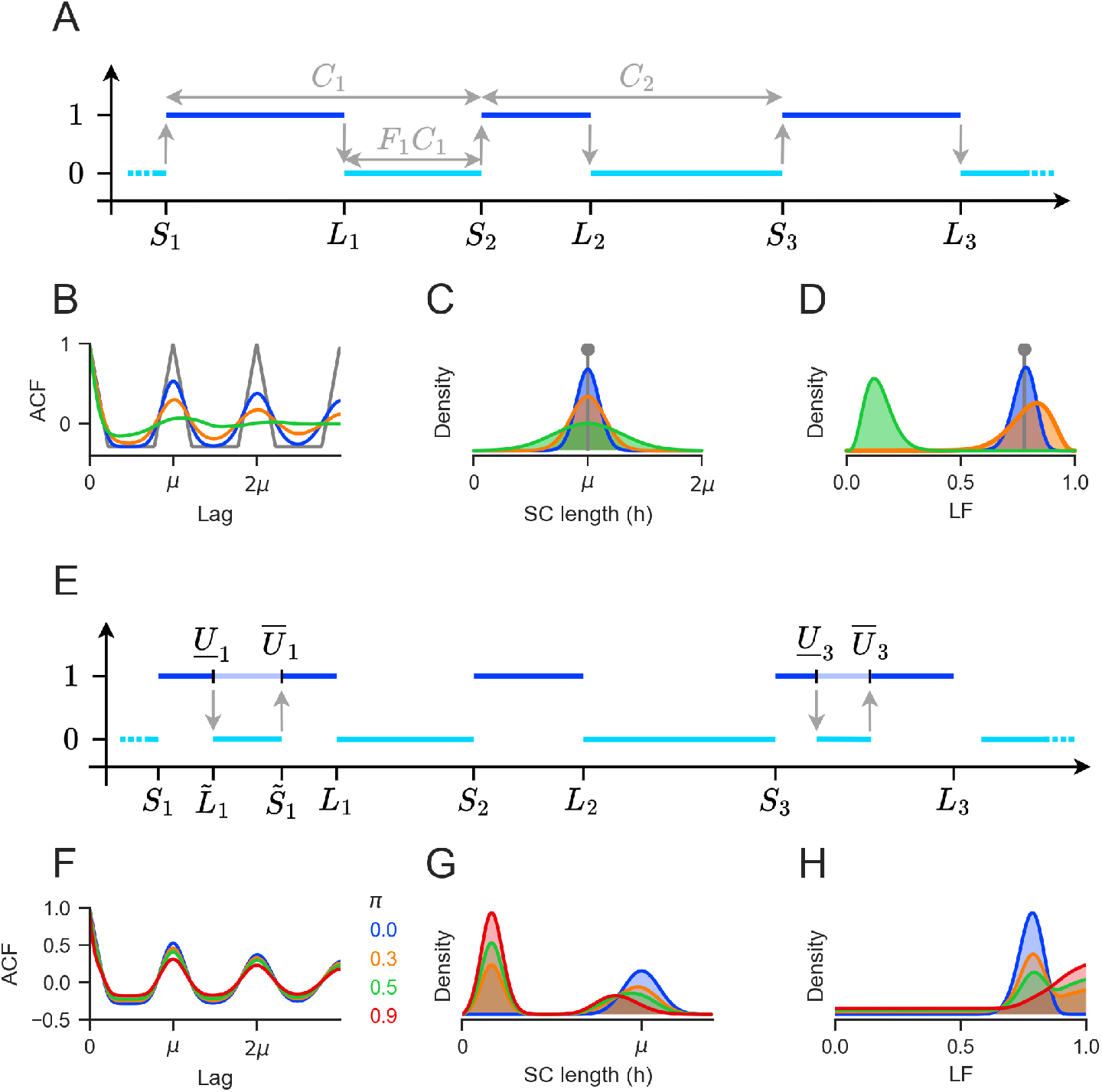
Model assumptions in the BN and IBN. **A**. The BN. SC lengths *C*_*i*_ between standing up *S*_*i*_ and standing up *S*_*i*+1_ are independent and truncated normally distributed. The lying fraction *F*_*i*_ between lying down *L*_*i*_ and standing up *S*_*i*+1_ of SC *i* is Beta distributed and independent. **B, C, D**. The ACF (B), the SC distribution (C) and the LF distribution (D) of a BN with three different parameter combinations (blue: *µ* = 100*m, σ* = 10*m, α* = 56, *β* = 16; orange: *µ* = 100*m, σ* = 15*m, α* = 16, *β* = 4; and green: *µ* = 100*m, σ* = 30*m, α* = 5, *β* = 30). **E**. The IBN. Given an underlying BN rhythm of standing and lying periods, each standing phase is interrupted independently with probability *π* in a random interval [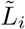, 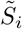], where the borders 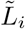, 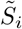 denote the minimum and maximum, respectively, of two independent Unif[*S*_*i*_, *L*_*i*_]-distributed random variables. Panel A shows a realization in which the first and third standing phase are interrupted. **F, G, H**. The ACF (F), the SC distribution (G) and the LF distribution (H) of a BN (blue) and IBN with interruption probability *π* = 0.3 (orange), *π* = 0.5 (green) and *π* = 0.9 (red).

To quantify regularity, we use the autocorrelation function (ACF). The ACF at lag *h* describes the correlation of a process with itself shifted by *h*. In this context, roughly speaking, it indicates the normed probability that an animal is standing at time *t* + *h*, given that it is standing at time *t*.

Figure 2 B illustrates various shapes of the ACF within a BN, along with the corresponding truncated normal (Figure 2 C) and Beta distributions (Figure 2 D). As a reference, we show in gray the ACF corresponding to a trivial deterministic model where all SCs have the same length *µ* and all lying phases have the same length. In this case, the ACF is periodic with period *µ* and shows perfect correlation at all multiples of *µ*. It then decreases linearly in both directions due to the linear decrease in expected overlap of the shifted standing phases with the original standing phases. When random variability in SC lengths and LFs is allowed, the ACF is dampened (Figure 2 B, blue, green, and orange curves corresponding to the SC and LF distributions in panels C and D). Thus, the height of the first side peak in the ACF serves as an indicator of regularity, taking the value 1 in the deterministic and perfectly rhythmic gray case and decreasing towards zero with increasing variability.

We now give the formal notation and definition of the BN and derive the formula for the ACF. Throughout the paper, we assume that standing and lying behavior in each night is described as a time-continuous stochastic process *X* which takes the value 1 during a standing phase and 0 during a lying phase (Figure 1 A). Random variables and processes are denoted by upper case letters throughout.

#### Definition 2.1 (The BN)

*Consider parameters µ* ∈ ℝ *and σ, α, β >* 0. *Let the process of standing up times S* := (*S*_*i*_)_*i*∈ℤ_ *be a stationary process with independent and identically truncated normally distributed increments S*_*i*+1_ − *S*_*i*_ =: *C*_*i*_ ∼ 𝒩 _+_(*µ, σ*^2^) *for i* ∈ ℤ, *where the subscript* _+_ *indicates the truncation at zero. Let further F*_*i*_ ∼ *Beta*(*α, β*) *be independent and beta distributed, and let*

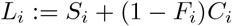

*denote the times of lying down. Then the Beta Normal (BN) process*

*BN*(*µ, σ*^2^, *α, β*) := (*X*_*t*_)_*t*∈ℝ_ *is defined as*

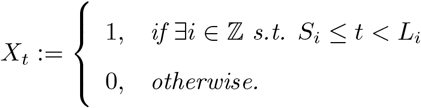

One night is assumed to be a finite interval of a realization of a BN *X* starting at a purely random time *t*_0_ ∈ ℝ.

The following theorem derives the expectation, variance and ACF of a BN process *X*. Because a BN (*X*_*t*_)_*t*∈ℝ_ is based on a stationary renewal process on ℝ of standing up times *S*_*i*_ with independent and truncated normally distributed intervals, *X* can be considered stationary. If we assume a night to start at a purely random time *t*_0_ ∈ ℝ, the distribution of (*X*_*t*_)_*t>t*0_ is identical for every *t*_0_ and thus, does not depend on *t*_0_, which implies stationarity also for a finite observed night (*X*_*t*_)_*t*∈[*t*0, *t*0 +*τ*]_, where *τ >* 0 denotes the length of the night. As *X* can only take the values 0 and 1, deriving the ACF reduces to finding the expected fraction of time during which the process is in the standing period both at a random time *t* and at time *t* + *h*.

#### Theorem 2.2 (ACF in the BN)

*Let X be a BN*(*µ, σ*^2^, *α, β*) *process as defined in Definition 2*.*1. Then the expectation and variance of X*_*t*_ *are given, respectively, by*

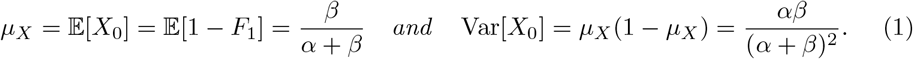

*The ACF of X is of the following form:*

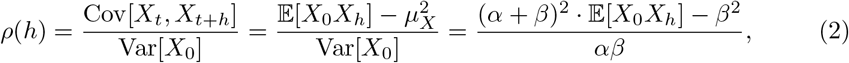

where

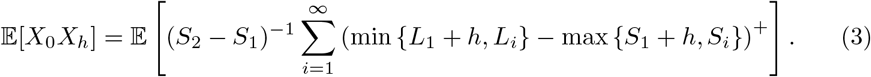

Equation (3) considers all combinations of standing phases in the SC overlapping *t* and all successive SCs because each SC has a standing period. Theorem 2.2 will turn out to be a special case of a stronger result in Theorem 2.5, hence its correctness will immediately follow from the proof of Theorem 2.5.

### 2.3 The BN with interrupted standing phases (IBN)

The BN is a parsimonious model for describing regular SC distributions. However, it fails to describe bimodal SC distributions characterized by many short SCs. One straightforward approach to model bimodal SC distributions would be to replace the normal distribution of SCs with a bimodal one, while maintaining the independence assumption of successive SC lengths. However, this would result in highly irregular processes that could not be coordinated with BN processes.

We therefore propose an extension of the BN that accounts for ultrashort SCs and bimodal SC distributions and that can directly be combined with a BN process, reproducing similar phenomena as shown in Figure 1 D. Specifically, we assume that the standing and lying behavior of animals with bimodal SC distributions is inherently no less regular than that of animals with unimodal SC distributions. Instead, bimodality of SC distributions may simply arise from a mechanism in which some standing periods are interrupted by short lying periods, such that short SCs could be observed in direct succession as in Figure 1 D (orange). This phenomenon is particularly expected in younger animals with a higher need for rest, who tended to exhibit the most prominent bimodal distributions with highest proportions of short cycles [21].

The BN with interrupted standing phases is called Interrupted Beta Normal (IBN, Figure 2 E). Interestingly, the size of the interruption probability *π* does not considerably change the regularity visible in the ACF (Figure 2 F), while the SC distribution is substantially transformed for larger interruption probabilities towards a bimodal shape (Figure 2 G), and also the LF distribution is affected (Figure 2 H).

We now give the formal definition of the IBN. In short, we assume a BN with rising times *S*_*i*_, *i* ∈ ℤ and times of lying down *L*_*i*_ as defined in Definition 2.1 in which each standing phase [*S*_*i*_, *L*_*i*_] is interrupted independently with probability *π* by a short lying phase 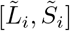 that starts and ends at two uniformly drawn time points within the standing period (Figure 2 E).

#### Definition 2.3 (The IBN)

*Let X be a BN*(*µ, σ*^2^, *α, β*) *process with standing cycles C*_*i*_, *lying fractions F*_*i*_, *standing times S*_*i*_ *and lying times L*_*i*_, *i* ∈ ℤ, *as given in Definition 2*.*1. Let π* ∈ [0, 1] *and let U*_*i*1_, *U*_*i*2_ ∼ Unif[0, 1] *and I*_*i*_ ∼ *Ber*(*π*) *for i* ∈ ℤ, *independent of C*_*i*_, *F*_*i*_, *S*_*i*_ *and each other. For each i* ∈ ℕ, *if I*_*i*_ = 1, *i*.*e. the standing phase is interrupted, define new times of lying down and standing up*, 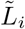 *and* 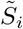, *respectively, as*

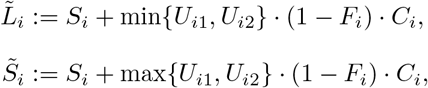

*and* 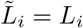 *and* 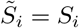 *if I*_*i*_ = 0. *Then the Interrupted Beta Normal (IBN) process IBN* 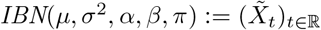 *is defined by:*

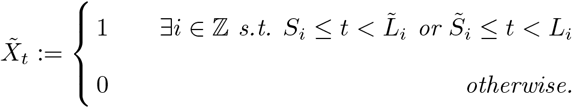

*The corresponding stand up times* 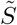 *of an IBN process are given by* 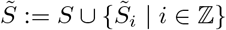.

Note that for *π* = 0, the IBN reduces to the BN.

The formal derivation of the ACF in the IBN is more complex than in the BN because each SC can exhibit two standing phases. Consequently, the infinite sum in equation (3) is expanded to four infinite sums. The ACF is provided subsequently by application of Theorem 2.5 to an individual IBN process.

### 2.4 Coordination across individuals: The synchronized IBN (S-IBN)

The IBN allows to investigate the degree to which two (I)BN processes synchronize their standing behavior (Figure 1 D). To that end, we assume that two processes *X* and *Y*, which can be BN or IBN processes, may share their underlying process *S* of standing up times. Note that in case of one or two processes being an IBN, we assume only the original standing up times *S* to be shared, not the standing up times 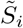 arising from interruptions of standing phases.

If *X* and *Y* share their standing up times *S* throughout the whole recording, we consider *X* and *Y* to be 100% coordinated. If, on the other hand, *X* and *Y* have two completely independent standing up processes *S*_*X*_, *S*_*Y*_, the processes would not be considered coordinated.

This synchronized model will be abbreviated S-IBN (Figure 3 A). Both the LFs and potentially interrupted standing phases are assumed to be drawn independently and with potentially different parameters (*α*_*j*_, *β*_*j*_, *π*_*j*_) for each animal *j*. The formal definition is given below in Definition 2.4.

**Fig 3.**
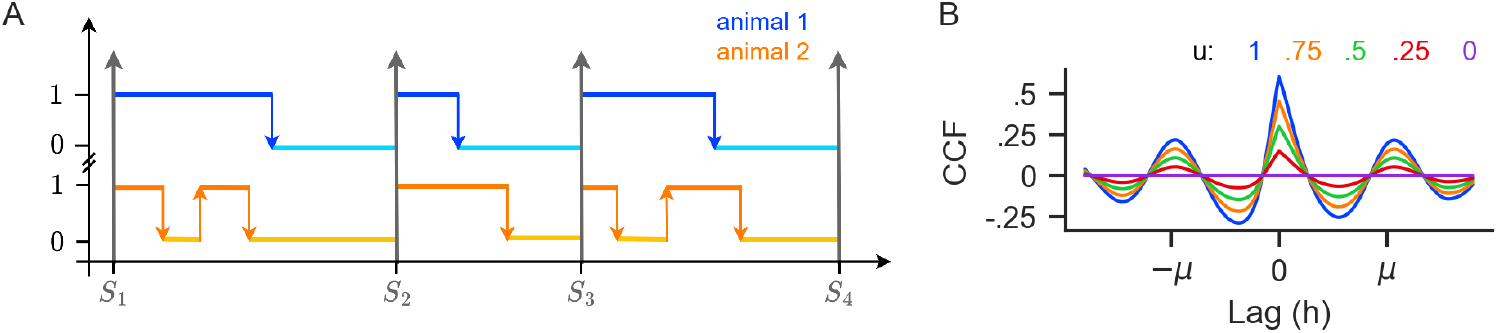
Coordination of standing behavior, the S-IBN and the crosscorrelation function (CCF). **A**. The S-IBN assumes that the standing up times *S*_*i*_ of two animals occur at the same time. The LFs are then drawn independently for the two animals (red and orange) from beta distributions with potentially different parameters. In cases of interrupted standing phases, interruptions are also assumed independent with potentially different interruption probabilities *π*_*j*_ for the two animals. **B**. Theoretical CCF in the IBN model (blue) with different values for the degree of coordination *u*.

Within the S-IBN, the standing up times of the two processes are perfectly synchronized. Under this assumption of perfect coordination, the crosscorrelation function (CCF) which is derived in Theorem 2.5, shows a joint oscillatory shape. Specific deviations from this shape can yield interesting biological insights into the degree of synchronization and potential temporal delays between the standing up times of the two individuals. These will be treated in Section 2.4.1.

We now give the formal definition of the S-IBN and the derivation of the CCF and ACF of IBN processes.

#### Definition 2.4 (The S-IBN)

*Let a stationary process of standing up times S be as defined in Definition 2*.*1 with independent and truncated normally distributed increments S*_*i*_ − *S*_*i*−1_ *for i* ∈ ℤ, *µ* ∈ ℝ, *σ >* 0. *Let the processes* 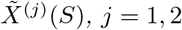, *be conditionally independent IBNs with parameters α*_*j*_, *β*_*j*_ *>* 0, *and π*_*j*_ ∈ [0, 1], *conditional on using the same process S of standing up times, i*.*e*., *with* 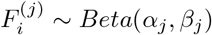, 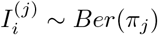 *and* 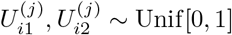 *for j* = 1, 2, *independent of C*_*i*_, *S*_*i*_ *and each other. Then the Synchronized IBN (S-IBN) process Z*_*S*_ *is defined by the two components*

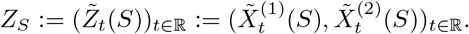

Theorem 2.5 includes the derivation of CCFs for all four cases in which one of the processes, both or none may show interrupted standing phases.

#### Theorem 2.5 (CCF in the S-IBN)

*Let Z*_*S*_ = (*X*^(1)^(*S*), *X*^(2)^(*S*)) *be a S-IBN process based on a joint process of standing up times S as defined in Definition 2*.*4 with parameters µ* ∈ ℝ, *σ, α*_1_, *β*_1_, *α*_2_, *β*_2_ *>* 0 *and π*_1_, *π*_2_ ∈ [0, 1]. *Then the CCF of Z*_*S*_ *is of the form*

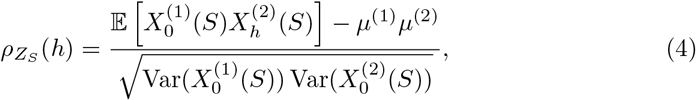

*where*

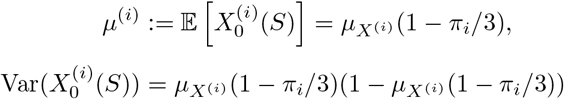

*and*

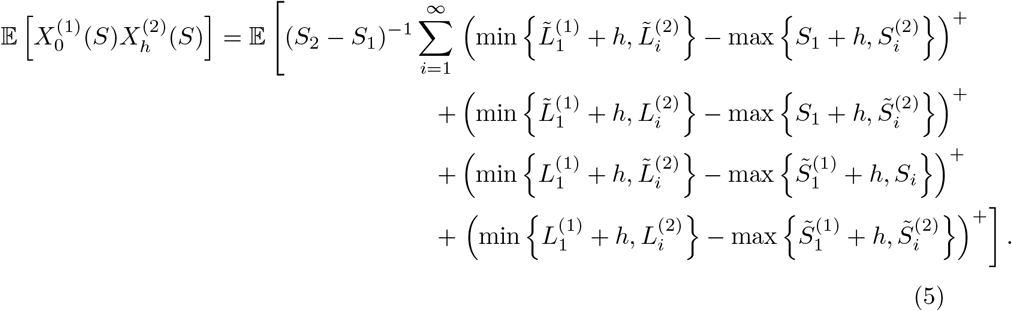

For the proof see Appendix 5.1. Calculating the CCF of a process with itself results in the ACF of the process. Therefore, the ACF of an IBN process 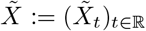 equals the CCF of the process 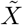 with itself, i.e., of 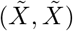 as given in Theorem 2.5.

Note that a further calculation of (5) is cumbersome, requiring infinite sums, multiple integrals and numerical integration. This holds irrespective of the assumptions on the underlying distribution. As a consequence, the ACFs and CCFs can be approximated faster by direct simulation of the terms 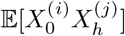.

#### 2.4.1 Temporal shift and degree of coordination in the S-IBN

As mentioned above, the shape of the CCF can give interesting biological insights concerning the degree of coordination or the temporal delay between standing up times. To investigate these issues, we introduce two further simple extensions of the S-IBN that will be used, first, to analyze potential temporal shifts between the standing up times of different animals, and second, to estimate the degree to which two processes share a common standing up rhythm *S*.

First, we assume an S-IBN process in which the standing up times of the second individual are shifted relative to the first by a delay *δ*. Formally, we assume a process 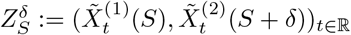 in which *S* + *δ* is defined by elements *S*_*i*_ + *δ, i* ∈ ℤ occurring at a deterministic delay *δ* after the standing up times in process 1. The CCF of 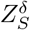 is then simply shifted by *δ* to the left, i.e.,

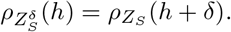

Second, we assume that the degree *u* to which two processes share the same standing up process *S* can vary continuously in [0, 1]. To that end, we assume that two IBN processes *X*^(1)^ and *X*^(2)^ share a background process *S* only for a fraction *u* ∈ [0, 1] of the recording period.

The degree of coordination *u* has a considerable effect on the CCF, dampening its oscillatory shape towards zero as *u* decreases because for *u* = 0, we find *ρ*_*Z*_ (*h*) = 0 for all *h*. Figure 3 illustrates theoretical CCFs of two IBN processes for different values of *u*. The formula for the CCF of two processes sharing the same background rhythm for a fraction *u* of the time is given in Lemma 2.6 (for a proof see Appendix 5.2).

##### Lemma 2.6

*Let Z* = (*X*^(1)^, *X*^(2)^) *be a S-IBN process with parameters µ, σ, α*_1_, *α*_2_, *β*_1_, *β*_2_ ∈ (0, ∞) *and π*_1_, *π*_2_ *as defined in Definition 2*.*4, and let S, S*_1_, *S*_2_ *be three independent processes of standing up times with parameters µ and σ. Let B* ∼ (*u*) *with probability u* ∈ [0, 1]. *Define*

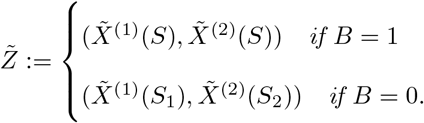

*Then the CCF of* 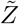 *is given by*

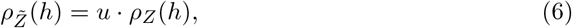

*with ρ*_*Z*_ *as in Theorem 2*.*5*.

### 2.5 Parameter estimation

The proposed (S-)IBN model can be used for several practical applications. First, on an individual basis, one can estimate the model parameters and analyse differences in, e.g., the oscillatory frequency *µ*, the regularity *σ/µ* or the interruption probability *π* as a function of the species, sex or age. Second, for pairs of individuals, one can investigate the degree of synchronization *u* or the delay *δ* between standing up times. We therefore provide here means for parameter estimation in the proposed models.

Parameter estimation for individuals within the (I)BN will be based on the raw SC and LF distributions, while estimation of *u* and *δ* for pairs of processes will be based on the CCF shape, which we compare to a CCF predicted under the assumption of *u* = 1 and *δ* = 0.

Note first that within the BN, the lengths of all observed SCs *C*_*i*_ and all LFs *F*_*i*_ are assumed independent. We could therefore estimate the parameters by maximization of the likelihood

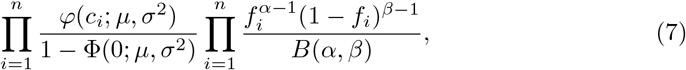

where *c*_1_, …, *c*_*n*_ denote the lengths of all fully observed SCs with associated LFs *f*_1_, …, *f*_*n*_. The first factor denotes the density of the truncated normal distribution, with *φ* and Φ the density and distribution function of the (untruncated) normal distribution, respectively. The second factor denotes the density of the Beta distribution. Both factors can be maximized numerically and individually.

For the IBN, however, parameter estimation is not as straightforward because the set of observable SCs also contains pairs of shorter SCs 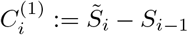 and 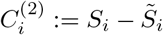, which result from splitting the respective original SC *C*_*i*_ if a standing phase is interrupted. The probability density functions (PDFs) of SCs and LFs in the IBN, *f*_*SC*_ and *f*_*LF*_, are therefore mixture distributions (see Appendix 5.3), and the lengths of successive observed SCs as well as their LFs are no longer independent. Maximum likelihood estimation would only be straightforward if we could distinguish between the original SCs and the SCs resulting from an interruption. As this is not possible, the likelihood in the IBN is extremely complex and would require the evaluation of all possibilities of interruptions between the observed standing phases.

For the IBN, we therefore estimate the parameters by fitting the theoretical PDFs of SCs and LFs to the empirical histograms of observed SCs and LFs (Figure 4 A-C, for details see Appendix 5.3). Figures 4 E and F show that the theoretical ACFs obtained with the parameter estimates should then match closely with the empirical ACFs.

**Fig 4.**
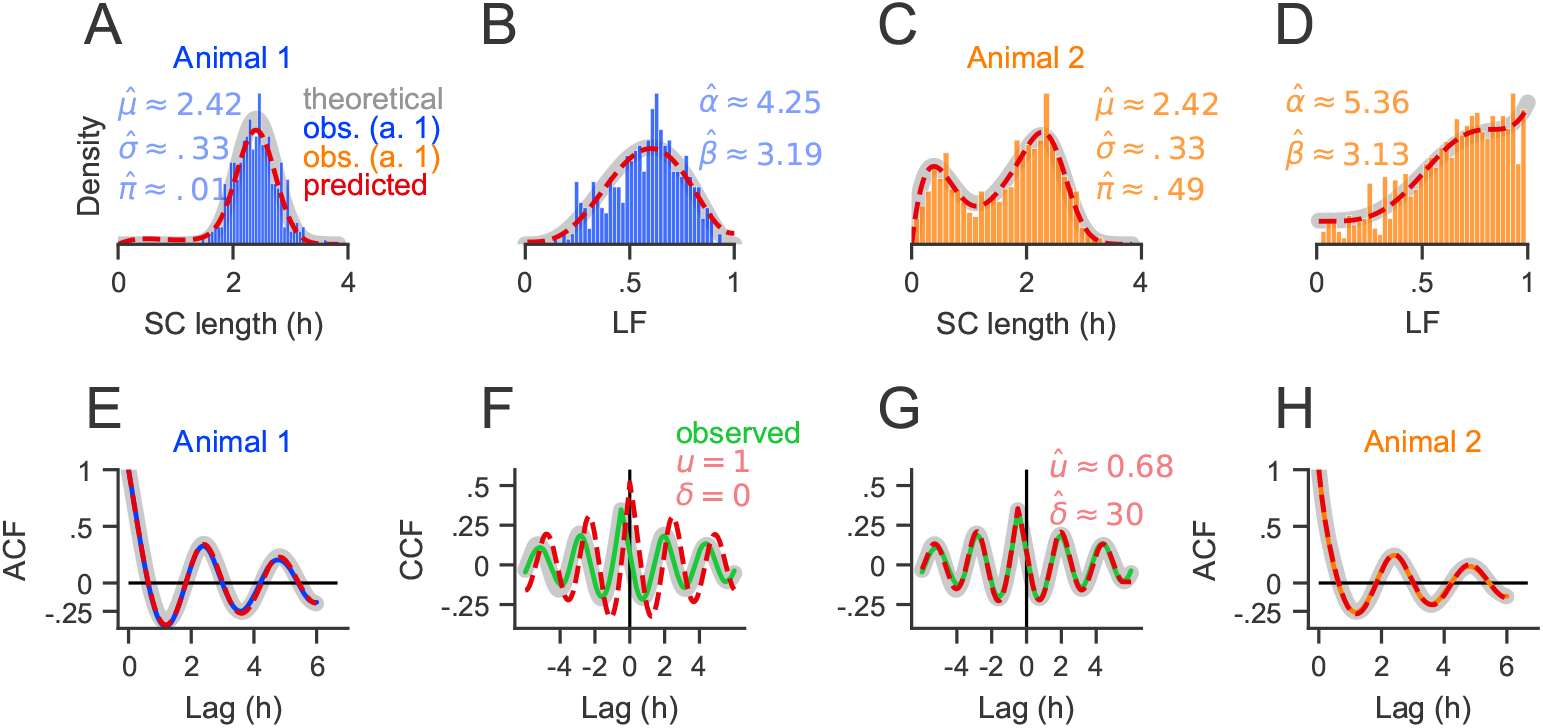
Top row: Theoretical densities (gray), simulated SC (blue in A/orange in C) and LF (blue in B/orange in D) distributions and fitted SC and LF densities (red) for two animals. Simulation parameters were *µ* = 2.4 hours, *σ* = 1*/*3*h, α*_1_ = 4, *β*_1_ = 3, *π*_1_ = 0, *α*_2_ = 5, *β*_2_ = 3, and *π*_2_ = 0.5, and about 450 and 900 simulated SCs, respectively. Bottom row: Theoretical (gray), simulated (blue/orange/green) and predicted (red) ACFs (E, H) and CCFs (F, G) for the two animals for the default parameters *u* = 1, *δ* = 0 in **F** and for the estimated parameters in **G**.

Note that the formulas for the PDFs of SCs and LFs (Theorem 5.1) reduce naturally to the respective densities in the BN for *π* = 0. We therefore propose to apply the IBN estimation for all processes because we cannot assume to know in practice in which case the BN holds, i.e., in which case *π* = 0.

An analogous fitting approach is applied for the S-IBN. Again, we first fit the theoretical PDFs of SCs and LFs to their respective histograms. Second, in order to estimate the parameters *δ* and *u*, we predict the CCF 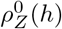 as in Theorem 2.5 with parameters 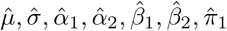, and 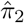 for an S-IBN with *u* = 1, *δ* = 0 (Figure 4 G) and then fit the function *u · ρ*_*Z*_(*h* + *δ*) to the observed CCF using nonlinear least squares (Figure 4 H).

#### 2.5.1 Estimation precision

In order to get an idea of the precision with which the parameters can be estimated and to evaluate the parameter estimates derived from the data set, we used simulations. To that end, we first extracted typical parameter combinations from the data set, which were used to simulate artificial nights. For each simulation, the parameters were then estimated and compared to the true parameters used in the simulation. For ease of notation, we do not indicate time units. Unless otherwise specified, all temporal parameters such as *µ, σ* and *δ* are measured in minutes, while *α, β* and *u* are dimensionless.

In order to identify typical parameters from the data, we first classified the SC distributions of the animals into unimodal and bimodal by visual inspection (cf. Section 3.2) and then chose *µ* = 144 as the approximate 75% quantile of mean SC lengths across all animals classified as unimodal for the simulations because the mean tends to underestimate *µ*, cf. Section 3.3. Similarly, we chose the 25% quantile of all standard deviations of SC lengths across all animals classified as unimodal, i.e., *σ* = 40. For the expectation of the LF, *α/*(*α* + *β*), we chose 0.68, the median of mean LF of unimodals, and for the standard deviation 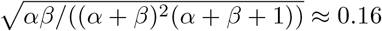, the mean standard deviation of LFs. The parameters *α* = 5.25 and *β* = 2.52 were chosen such as to obtain similar expectation and standard deviation of LFs. In addition, we chose different interruption probabilities *π* ∈ {0, 0.3, 0.5, 0.7, 0.9}. Synthetic data were then generated according to the IBN for different numbers of nights, and then fitted for each combination of parameters.

Figure 5 shows that the estimation error decreases with the number of nights, where small errors below about 10% are obtained for most parameters with 50 or more recorded nights (Fig. 5 A-D). Simulations also indicate that in the BN, i.e., if *π* = 0, estimates of *π* are close to zero. One should, however, also recall that in the present data set, very short standing and lying periods shorter than 5 minutes had been eliminated in order to reduce automatic misclassification of standing or lying behavior. We therefore repeated the simulations, while also eliminating such short standing or lying phases. Figure 5 E-H indicate that this preprocessing may increase estimation error, particularly for large interruption probabilities *π*. This is plausible because the particular bimodal shape of the SC distribution is systematically distorted by elimination of very short cycles, such that particularly positive values of *π* will tend be underestimated (Fig. 5 H).

**Fig 5.**
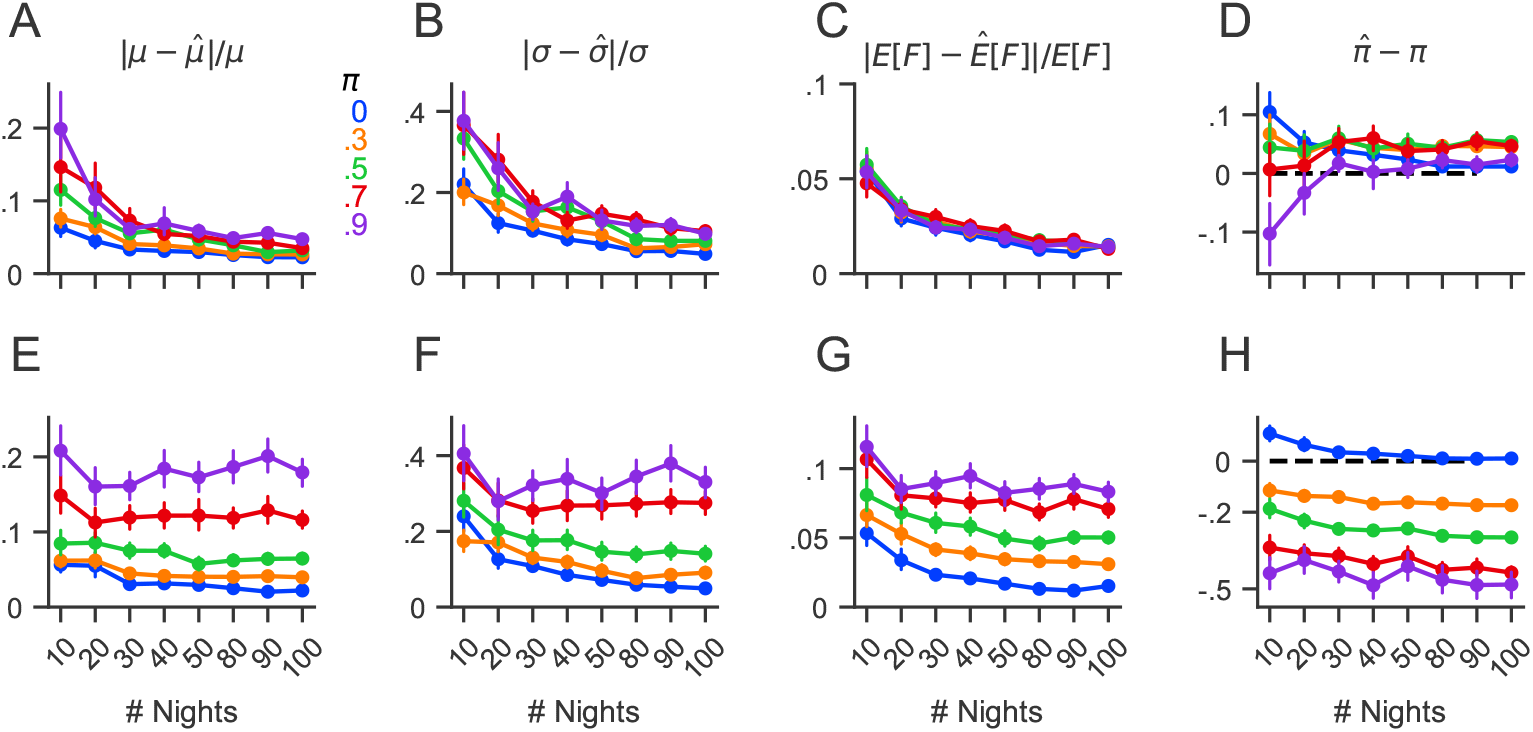
Estimation precision in the IBN with 40 bins for simulated data with *µ* = 144, *σ* = 40, *α* = 5.25 and *β* = 2.52 and *π* ∈ {0, 0.3, 0.5, 0.7, 0.9}, 100 simulations per parameter set. Top row: Mean relative error of **A**. *µ*, **B**. *σ*, **C**. 𝔼[*F*], and error of **D**. *π*, with error bars indicating standard error. Bottom row: **E, F, G, H**. Analogous simulations in which additionally, as performed in the preprocessing of the present data set, lying and standing phases shorter than five minutes were removed as described in [21].

In order to investigate the estimation precision of the degree of coordination *u* and the delay *δ* in the S-IBN, we simulated pairs of processes according to the S-IBN model as specified in Section 2.4 for different numbers of nights and then fitted the S-IBN for each combination of parameters. In the simulated parameter range, the estimation error of 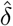 was rather small relative to the period *µ* and decreased with the number of nights. Because estimates of *u* are restricted to the unit interval [0, 1], *u* tended to be slightly overestimated for *u* = 0 and underestimated for large values of *u*, where again the estimation error decreased with the number of recorded nights.

## 3 Results

### 3.1 Data description

In the present analysis, we included all animals of the data set reported in [21] that showed at least 30 completely observed SCs. As a result, three adult animals (one male *Oryx dammah* and two female *Equus quagga*) were excluded from the data set, resulting in 189 animals (60 male, 129 female) representing 19 taxa from 20 different zoos. A median number of 47 nights was observed per individual, ranging from 11 to 185 nights per individual. Out of the 189 observed animals, 141 spent all nights alone in a box, while 40 animals consistently shared their box with the same companion every night. Seven further animals shared their box only in a small subset of nights with one companion, where the companion also sometimes changes. For one particular animal (a female adult *O. dammah*), we could observe a long period with the same companion, and then another long period with a different companion (see Section 3.7).

Out of all 189 animals, 16 were classified as young, 20 as subadults and 153 as adult as in [21]. Among the 21 consistent pairings, most (14) were adult-young pairs, 3 were adult-adult pairs, two were subadult-young pairs, and one was subadult-subadult and adult-subadult, respectively.

### 3.2 Motivation of the model assumptions

#### BN assumptions

The model assumptions were largely motivated by empirical observations. First, for many individuals, the SC lengths appeared to be roughly normally distributed (see Figure 1 B and also [21] for the corresponding distribution of the lying cycles). Indeed, a subset of 150 out of 189 individuals was identified by visual inspection showing a roughly unimodal and symmetric SC distribution suitable for a description in the BN. These individuals were included in the investigation of the further BN model assumptions.

Second, we observed considerable differences between the SC distributions of different individuals even of the same species, sex and age group (Figure 7 A), motivating the assumption that parameters *µ*_*j*_ and *σ*_*j*_ of the SC distribution may depend on the specific individual *j*. Third, serial correlations between SC lengths was rather small with a mean of 0.06 (*±* SEM of 0.0995), motivating the assumption of independence of SC lengths. Fourth, the assumption of identically distributed SC lengths was motivated by the observation that only weak dependency of the SC length on the particular time during the night was observed. To a rather small extent, the SC lengths sometimes increased minimally across the night, in the mean by about 2.3 minutes per hour across all animals (*±* SEM of 11.8 seconds, Figure 7 C). As this effect was rather small, we assumed identical distribution, which largely simplifies technical derivations.

**Fig 6.**
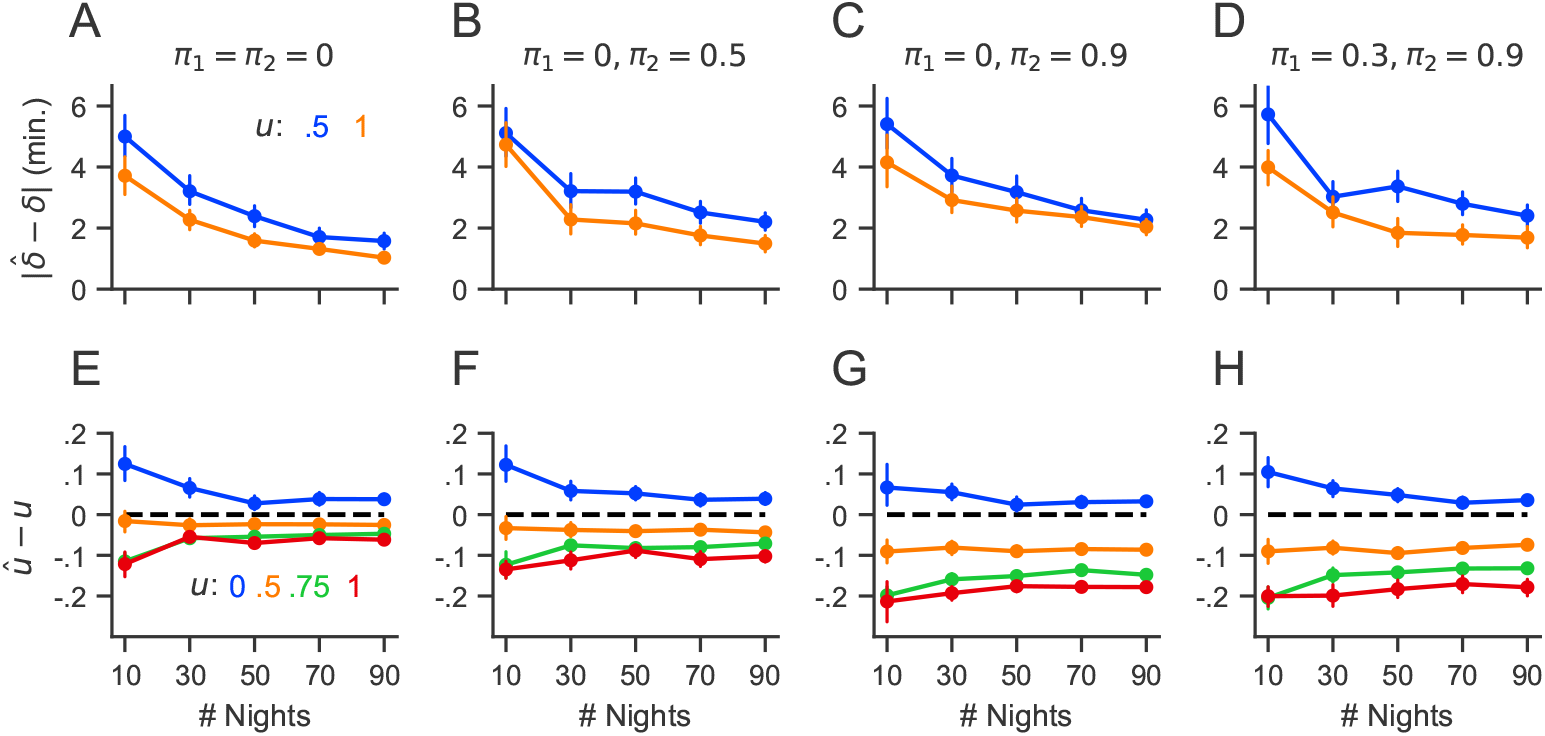
Estimation precision in the S-IBN with 20 bins for simulated data with *µ* = 144, *σ* = 40, *α* = 5.25, *β* = 2.52, (*π*_1_, *π*_2_) and *u* as indicated, *δ* = 0. Simulations were performed with elimination of short standing or lying periods. Error bars indicate standard errors. Top row: Mean absolute error of 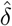 for different values of (*π*_1_, *π*_2_) and *u* as indicated. Bottom row: Mean error of *û* for *π*_1_ = 0 & *π*_2_ = 0 (E), *π*_1_ = 0 & *π*_2_ = 0.5 (F), *π*_1_ = 0 & *π*_2_ = 0.9 (G), and *π*_1_ = 0.3 & *π*_2_ = 0.9 (H), and *u* as indicated.

**Fig 7.**
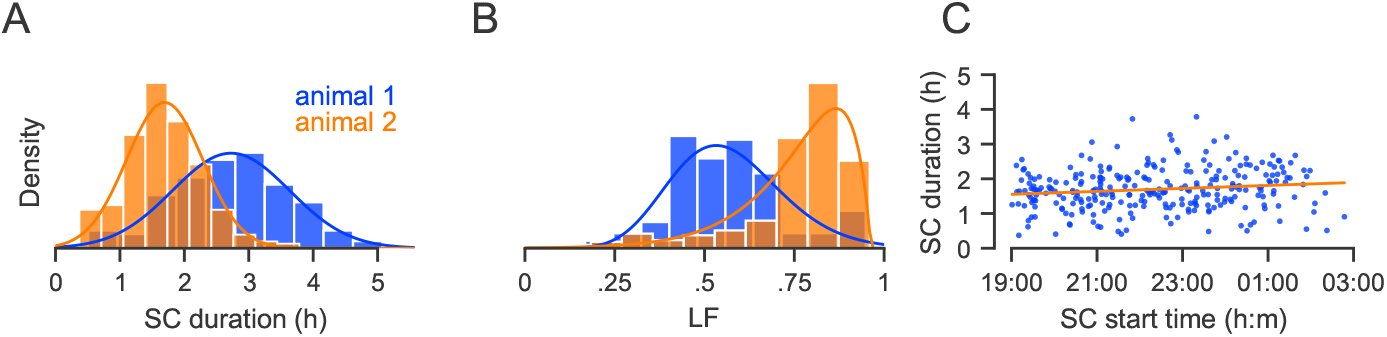
Motivation of BN model assumptions. (A) Histograms of SC length distributions in two adult female *C. taurinus* and fitted truncated normal distributions as described for the BN in Section 2.5. (B) Empirical LF distributions and fitted Beta distribution densities for the two animals in A. (C) SC duration as a function of SC start time for a female adult *Oryx leucoryx*, with standard linear regression line (orange).

Finally, also the lying fraction did not show a considerable serial dependence (−0.05 *±* 0.012, mean *±* SEM of Pearson correlation coefficient) nor any considerable dependence on the length of the respective SC (0.1 *±* 0.03, mean *±* SEM of Pearson correlation coefficient), motivating independence of LFs. The Beta distribution was considered sufficiently flexible to describe various LF distributions observable in different animals (Figure 7 B).

#### IBN assumptions

The nocturnal behavior of animals classified visually as unimodal could often be described well with the BN (Figure 8 A-D), where real and simulated nights showed high correspondence, and also the shape of the ACF could be predicted from the BN fit (Figure 8 I). For the bimodal case, however, this was often not the case (Figure 8 E-G). Specifically, bimodality could not be captured by the truncated normal distribution (red curve in Figure 8 E), simulated nights showed less regularity and fewer small SCs (Figure 8 G) than observed empirically (Figure 8 H), and the empirical ACF showed a more regular shape than predicted from the BN assumptions (Figure 8 J).

**Fig 8.**
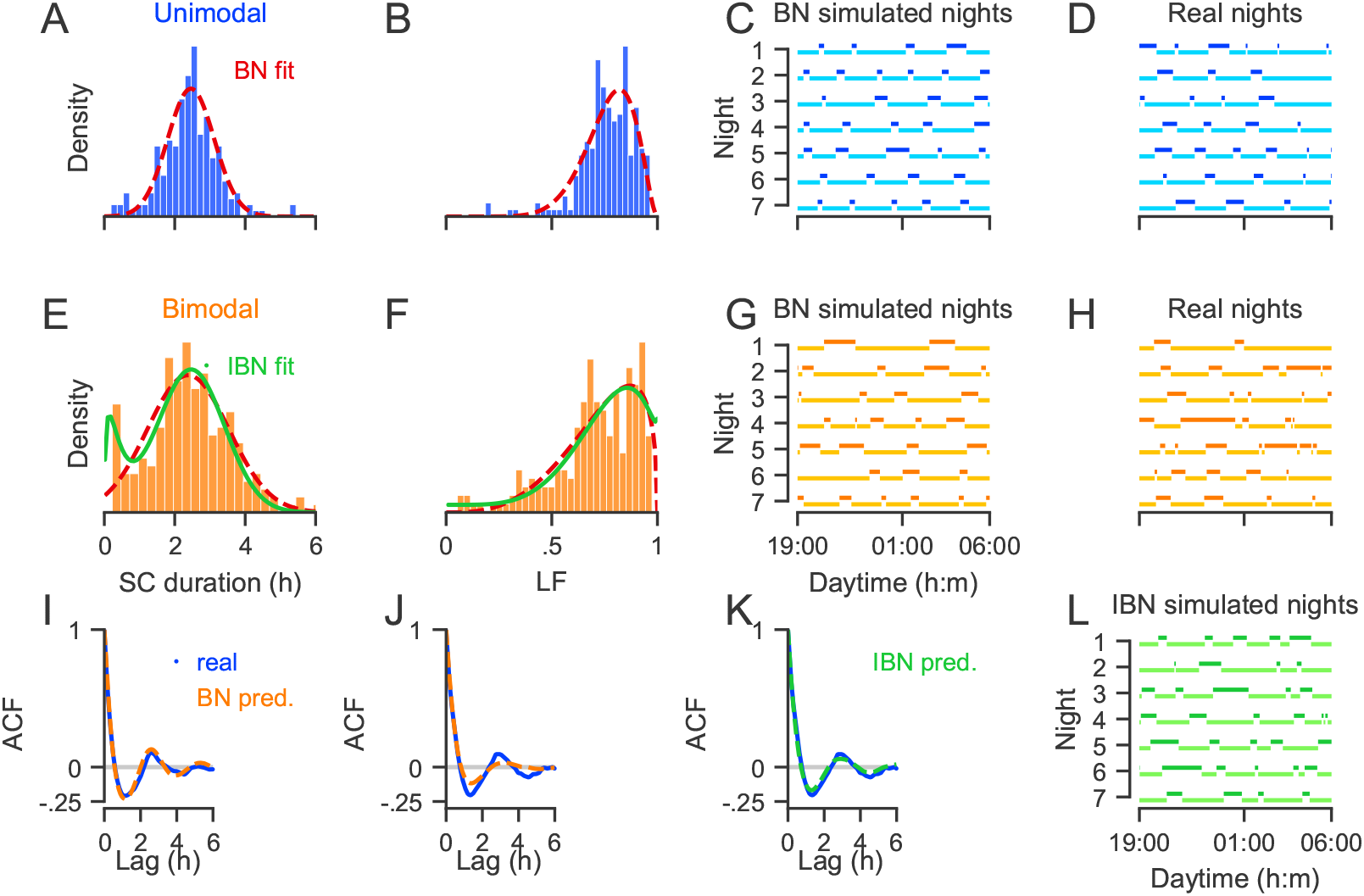
Motivation of IBN model assumptions. A-D: BN model fitted to the SC distribution (A) and LF distribution (B) of an animal with unimodal SC distribution. Simulated BN nights (C) and observed nights (D) showed high correspondence, as well as empirical and predicted ACFs (I). E-H: BN model (red) and IBN model (blue) fitted to the SC distribution (E) and LF distribution (F) of an animal with bimodal SC distribution. Low correspondence between nights simulated with BN assumptions (G) and empirical nights (H) and between empirical ACF and ACF predicted with BN assumptions (J). (K) ACF for bimodal animal predicted with IBN assumptions, and simulated nights under IBN issumptions (L) show close agreement to empirical observations.

By introducing only the interruption probability *π* as one additional parameter, the IBN could often capture these features, enabling a better fit of the bimodal distribution (Figure 8 E, green), a more suitable, regular shape of the ACF (Figure 8 K) and a closer description of empirical nights (Figure 8 L).

In the next section, we will use the insights of the IBN to analyse the seemingly irregular behavior of juvenile individuals.

### 3.3 No significant difference in IBN based regularity measure

We first fit the whole data set individually, applying the IBN to all individuals as described in Section 2.5. The parameter estimates of all animals are presented in supplementary Figure S1. Consistent with the findings in [21], no considerable differences could be observed in the SC parameters *µ* or *σ* as a function of species, age or sex, but zebras usually showed much lower LFs than bovids.

Note two differences between the present analysis and [21]: First, the analysis in [21] was applied to periods between two lying cycles, i.e., two successive events of lying down, while the present analysis was applied to SCs, i.e., periods between two successive events of standing up. This, however, should not largely affect the interpretation. Second, and more importantly, [21] compared the raw mean and standard deviation, *m* and *s*, while we compare the estimates of *µ* and *σ* of the underlying, truncated and potentially interrupted normal distribution in the theoretical IBN. The raw mean *m* is similar to the parameter *µ* but tends to underestimate *µ*, especially in the IBN because the interruption of SCs yields a bimodal SC distribution with an empirical mean that is systematically decreased relative to *µ*. In a similar way, the raw standard deviation *s* is similar to *σ* but tends to overestimate *σ* in the IBN because it is based on the whole mixture distribution including also the interrupted, short SCs.

As a consequence, the standard coefficient of variation (CV), *s/m*, tends to underestimate the regularity. Instead, we propose to use the IBN-CV, 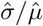, of the underlying SC rhythm, which should be smaller than the CV because it reflects the base rhythm of the individuals. Indeed, this effect could also be observed in the dataset, where the IBN-CV was typically smaller than the standard CV (Figure 9 A, *p <* 0.0001, Wilcoxon test).

**Fig 9.**
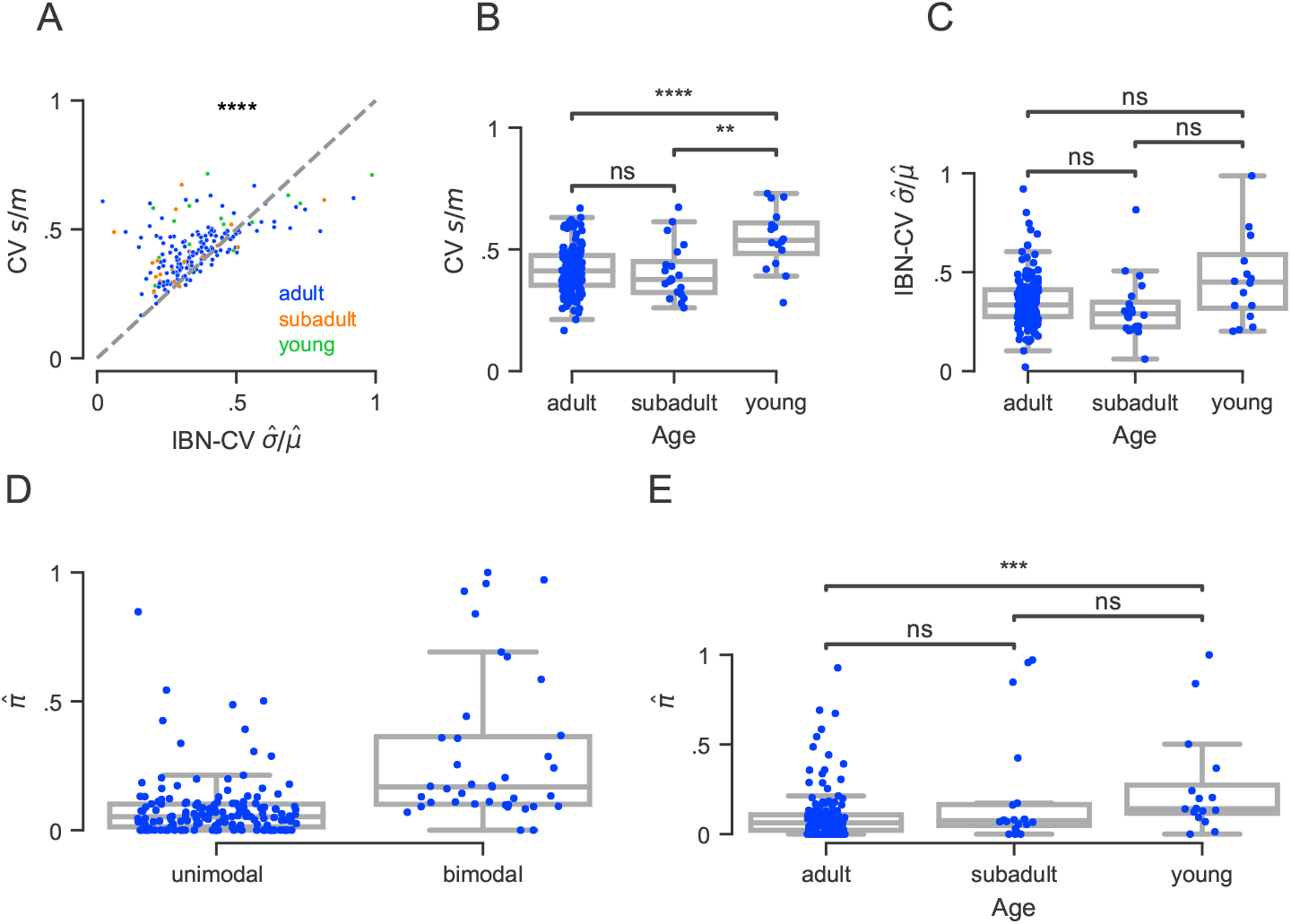
Comparison of regularity and interruption probability across age groups. **A**. Comparison of the difference in the raw CV, *s/m*, and the IBN-CV 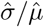, where 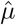 and 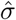 are obtained by fitting the IBN to the observed standing cycles. **B**. CV and **C**. IBN-CV as a function of age group. **D**. Estimates of the interruption probability *π* for animals initially classified as unimodal and bimodal. **E**. Estimates of the interruption probability *π* as a function of age group. Stars indicate significance levels (n.s. *p >* 0.05, * *p <* 0.05, ** *p <* 0.01, *** *p <* 0.001, **** *p <* 0.0001).

With the new IBN, we thus observed differences in the interpretation of the observed regularity: On the one hand, similar to the results reported in [21], the standard CV showed significant differences between adult and young animals (Figure 9 B *p <* 0.001) and between subadult and young animals (*p <* 0.01), indicating a seemingly higher irregularity in younger animals. On the other hand, the differences in the IBN-CV were no longer significant between adult and young animals (*p* = 0.11), between adult and subadult animals (*p* = 0.07), and between subadult and young animals (*p* = 0.06), i.e., the hypothesis that the regularity that underlies the SC rhythm is comparable for adult and young animals could no longer be rejected (Figure 9 C, *p* = 0.1).

### 3.4 Higher interruption probabilities in younger animals

While the IBN-CV, i.e., the regularity of the underlying rhythm, showed no significant differences between age groups, younger animals showed higher interruption probabilities than adult animals (Figure 9 E, *p <* 0.001, adult vs. subadult: *p* = 0.35, subadult vs. young: *p* = 0.09). Out of 16 young animals, 14 (87.5%) showed an estimated interruption probability of 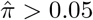, as compared to 15/20 (75%) in subadult and 84/153 (54.9%) in adult animals (*p* = 0.014, Chisquare Test).

The estimated interruption probabilities were also in high agreement with the visual bimodality classification performed in Section 3.2: Out of 39 animals classified as bimodal, 37 had a low interruption probability close to zero, i.e., 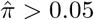, whereas interruption probabilities for animals classified as unimodal were lower, almost 50% of unimodals showing 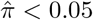 (Figure 9 E).

Thus, in the IBN, age specific differences were not found in the regularity of the underlying rhythm but in the increased tendency to interrupt standing phases by short lying phases in younger individuals.

### 3.5 Degree of coordination decreases with spatial distance

We then estimated the degree *u* to which pairs of animals coordinated their standing and lying activity, as described in Section 2.4. To that end, we applied the fitting approach described in Section 2.5 to all pairs of animals that were sharing the same box (i.e., a box distance of zero). In addition, we investigated a random subset of pairs of animals that were standing in adjacent boxes in the same building, i.e., with box distance one, and other pairs of animals that were standing at least two boxes apart from each other, in the same building.

Figure 10 A shows a decrease in the estimated degree of synchronization *û* of the animals’ underlying SC rhythm, from a mean of 0.8 (SEM 0.04) for pairings in the same box to 0.26 (SEM 0.03) in adjacent boxes, and 0.12 (SEM 0.03) in boxes further apart. Differences were significant between same-box and adjacent-box pairings (*p <* 0.00001, Wilcoxon test), but not significant between adjacent boxes and boxes further apart (*p* = 0.13). Figures 10 D-F show typical examples of empirical and fitted CCFs for animal pairs with different box distances, illustrating the decrease in coordination with increasing distance.

**Fig 10.**
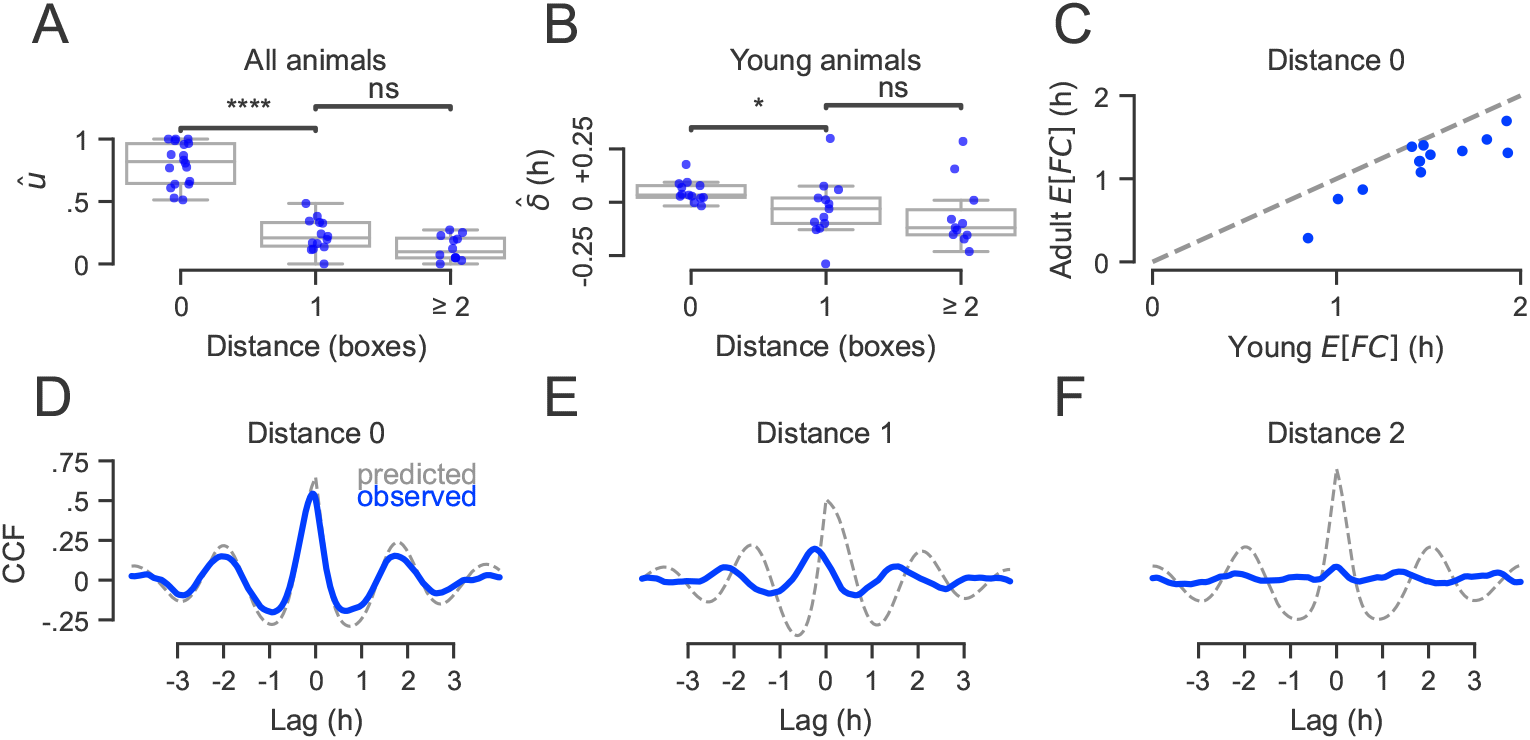
**A**. Fitted values of *û* by box distance for all pairs of animals of same taxa. **B**. Fitted values of 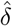 by box distance for all pairs of animals involving a young animal. **C**. Comparison of the estimated expected lying time of young animals and their adult box partner. **D**. Predicted (gray, *u* = 1) and observed (blue) CCF for one young female *Tragelaphus Oryx* sharing a box with a subadult female *T. Oryx*. **E**. Predicted (gray, *u* = 1) and observed (blue) CCF for the young animal from (D) with a randomly chosen adult female *T. Oryx* from a neighboring box (distance of 1). **F**. Predicted (gray, *u* = 1) and observed (blue) CCF for the young animal from (D) with a randomly chosen adult male *T. Oryx* with a distance of two boxes.

### 3.6 Young animals tend to stand up later and lie down earlier

In order to investigate the temporal delay *δ* between the standing up times of the animals, we focused only on pairings with young animals. This is because for any pairing of animals AB, it holds that *δ*_*AB*_ = −*δ*_*BA*_ and thus, the sign of the estimated delay is arbitrary if the two animals belong to the same group. We therefore estimated only the delays *δ*_*AB*_ for which animal A was a young animal (Figure 10 C). Interestingly, almost all delays were positive (mean delay of 181.77 s, SEM 50.65 s) for pairings in the same box, indicating that relative to the standing up time of the adult animal, the paired young animal would be getting up on average 3 minutes later. Delays for pairings with distance of one or more boxes were not significantly different from zero (*p* = 0.9 for box distance one and *p* = 0.46 for box distances greater than one). Consistently, differences of delays between a box distance of zero and one were statistically significant (*p <* 0.05, paired Wilcoxon test).

Interestingly, one can also combine the delay time *δ* of standing up in young animals with their expected lying duration within a cycle. Recall that *C* denotes a cycle length and *F* the lying fraction, such that 𝔼(*FC*) denotes the expected lying duration within a cycle. This expected duration was longer in young animals (Figure 10 C, *p <* 0.001, Wilcoxon test). From these we can estimate, for each individual, the average time that the young animal would *lie down* in relation to the lying time of the paired adult animal. Typically, young animals would lie down in the mean 17.6 minutes (SEM 2.76 m) before the adult.

### 3.7 Changes in individual rhythms – a case study

In one particular pairing of animals, an interesting change of pairings was observed during the recording period. One female adult *O. dammah* first shared a box with a subadult male *O. dammah* for 113 nights, then spent three nights alone before sharing a box with a young male *O. dammah* for 69 nights. The originally paired subadult then spent 61 nights alone in a box.

In order to compare the rhythms and SC distributions of this animal across the different pairing conditions, we analysed, first, the raw, i.e., model free, SC distribution. Second, we fitted the S-IBN to to the different pairings to investigate differences in the degree of synchronization.

In the raw SC distribution, we observed a striking change for the same adult animal (Figure 11 A, blue). In the nights spent with the subadult, the mean SC length of the adult was 115.7 min (SEM 2.82 min), while the mean SC length of the adult was much higher (149 min, SEM 4.04 min) during the nights spent with the young animal (*p <* 0.0001, Wilcoxon test). The mean SC of the young animal (141.03 min, SEM 6.21 min, Figure 11 A, orange) was much higher than the mean SC of the subadult animal (109.58 min, SEM 3.71 min) when standing alone (Figure 11 A, green), this difference being highly significant (*p <* 0.0001). In comparison, no significant difference in SC lengths could be observed in the subadult animal between standing alone and being paired with the adult (*p* = 0.94), and no significant difference could be observed between the SC lengths of the adult and subadult when sharing a box (*p* = 0.18), as well as between the SC lengths of adult and young animal when sharing a box (*p* = 0.09). This suggests that the adult animal may have adapted its SC rhythm when the pairing changed, while no adaptation could be observed in the subadult animal.

**Fig 11.**
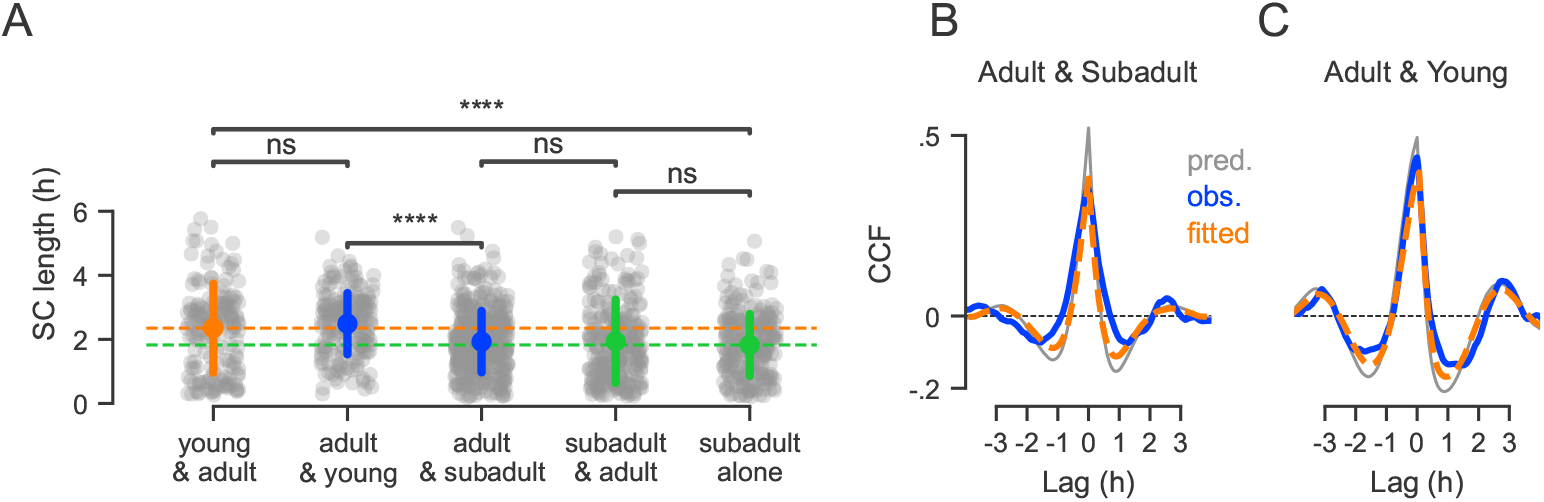
**A**. Comparison of SC lengths between different pairings. Points and bars indicate mean and standard deviation, respectively. Horizontal lines indicate mean SC lengths of the young animal when paired woth the adult (orange) and of the subadult animal (green) **B**. Observed (blue), predicted (gray, with *u* = 1) and fitted (orange, with *û* = 0.74) CCF of adult and subadult animal. **C**. Observed (blue), predicted (gray, with *u* = 1) and fitted (orange, with *û* = 0.81) CCF of adult with young animal, exposing higher *û* than in (B).

In addition, in the S-IBN based analysis, we observed a considerable change of the degree of coordination within the adult animal. While coordination was moderate when paired with the subadult animal (*û* = 0.74, Figure 11 B), coordination was enhanced when paired with the young animal (*û* = 0.81, Figure 11 C).

## 4 Discussion

In this paper we have analysed the nocturnal standing and lying activity of a large number of animals belonging to different ungulates, and between 11 and 185 nights were observed for each individual.

Our first aim was to investigate regular as well as seemingly irregular behavior, particularly such behavior with remarkably many short standing phases as often observed in young animals. To that end, we have proposed a natural stochastic model called IBN that captures both, the regularity and the existence of short SCs with a few simple assumptions. Most importantly, it assumes that a regular rhythm persists, but that standing phases within this rhythm may be interrupted by short lying phases. In this way, very short standing phases arise in an otherwise regular rhythm.

Using this model for parameter estimation of the underlying rhythm and the ACF for quantification of regularity allows to reveal a higher regularity than would be visible only from the raw distributions of SCs. Application to our recordings showed no significant differences in our new regularity measures between the age groups, while in contrast, the probability that a standing phase was interrupted by a short lying phase was increased in younger animals. This suggests that seeming irregularity may in fact not necessarily reflect a reduced background regularity, but instead, a higher probability that standing phases are interrupted by short lying phases. In this context, one should also keep in mind that the interruption probability, if positive, tends to be underestimated when applying a preprocessing in which ultrashort standing or lying phases are eliminated prior to the analysis. This suggests that interruption probabilities might be even higher than estimated in the present data set.

Secondly, we investigated the coordination of standing and lying for pairs of animals that were stalled close to each other. To that end, the IBN was naturally extended by assuming that a certain fraction *u* of the rhythm of standing up times occurred in a synchronized manner (or potentially with a fixed delay *δ*), for the two animals. A high degree of coordination up to 100% was observed in animals sharing the same stable box, where this coordination decreased strongly with the distance between boxes. Young animals sharing a box with an adult animal tended to stand up about 3 minutes after the adult and lie down earlier.

Thus, our findings suggest that all recorded animals showed a more or less regular underlying SC rhythm, where younger animals tended to stand up in a synchronized but slightly delayed manner in relation to the adult, then sometimes interrupting their standing phase by shortly lying down and standing up again, and then lying down again before the adult animal.

In an additional case study, we even observed a significant change in individual SC rhythm of one adult individual when changing its partner from a subadult to a young animal as well as a change in the degree of coordination.

The model assumptions within the IBN are simple and highly motivated by empirical observations. Interestingly, the model was applicable to all observed species, suggesting that all animals showed a certain regularity in their standing-lying behavior. In principle, a generalization to other comparable time series that describe switching between two states is possible as long as the switching occurs with a certain underlying regularity.

Technically, in spite of the simplicity of the model assumptions, parameter estimates cannot be obtained in closed form, and also ACFs and CCFs need to be derived numerically. This restricts statistical possibilities for, e.g., individual confidence intervals or comparisons of individual parameter estimates but represents less of a problem when many nights or individuals can be observed.

In similar models for point processes, the derivation of ACFs and CCFs is more straightforward [25–27] and might be an interesting development for future studies if one intends to focus only on the times of standing up or on the times of lying down. Also, an extension to more than two processes would be highly interesting, for example in cases in which more than one or two individuals are stalled together in the same enclosure.

In summary, we believe that the present model can provide interesting new insights into regular nocturnal standing and lying behavior and similar time series.

## Acknowledgments

This work was supported by the LOEWE Schwerpunkt CMMS – Multiscale modelling in the life sciences and received financial support from von Opel Hessische Zoostiftung. We would like to thank Götz Kersting for providing an intuitive argument for the stationarity of the introduced processes.

## 5 Appendix

### 5.1 Proof of Theorem 2.5

As the two processes *X*^(1)^ := *X*^(1)^(*S*) and *X*^(2)^ := *X*^(2)^(*S*) of a S-IBN process *Z*_*S*_ = (*X*^(1)^(*S*), *X*^(2)^(*S*)) are jointly stationary, their cross correlation function (CCF) at lag *h* ∈ ℝ is given by

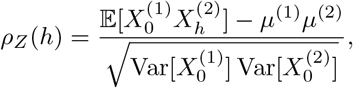

with 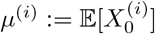.

In order to derive 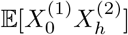, we need to consider several cases. W.l.o.g. we assume that 0 ∈ [*S*_1_, *S*_2_). Since *S* is stationary, the concrete position of zero in the interval [*S*_1_, *S*_2_) is uniform, i.e. (0 − *S*_1_)*/C*_1_ ∼ Unif[0, 1]. Thus, given 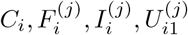 and 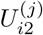 for *i* ∈ ℤ and *j* = 1, 2, computing 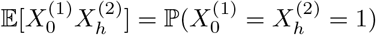 reduces to measuring the overlap of standing phases, i.e., phases during which both processes 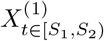 and 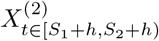 equal 1, normalized by *C*_1_ = (*S*_2_ − *S*_1_). Since the standing phases are given by 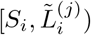 and 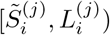 for *j* = 1, 2, this is equivalent to measuring the overlap for all four combinations of standing phases. The overlap of two intervals [*a, b*) and [*c, d*) is given by (min{*b, d*} − max{*a, c*})^+^. Hence we need to derive

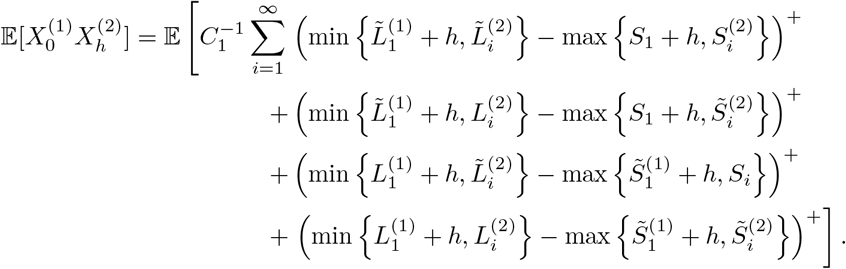

Reducing the CCF to the autocorrelation function (ACF) of a process *X* results from setting *X*^(1)^ = *X*^(2)^ =: *X*. In the BN case in Theorem 2.2, only one standing phase is observed per standing cycle, resulting in only one infinite sum for 𝔼[*X*_0_*X*_*h*_] as given in equation (3). It remains to derive *µ*^(*i*)^ and Var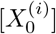, which we do for the general IBN case. The BN case (Theorem 2.2) follows with *π* = 0. For an IBN process 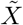, the expectation and variance can be derived as

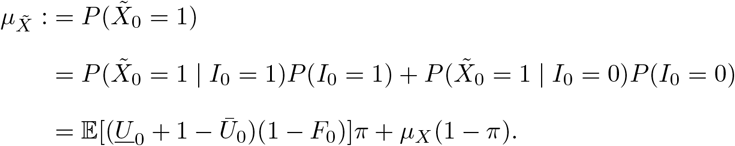

Due to independence of (*<u>U</u>*_0_, *Ū*_0_) and *F*_0_ and using 𝔼*Ū* = 2*/*3 and 𝔼*<u>U</u>* = 1*/*3 (see Appendix 5.2), we get

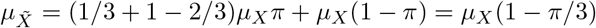

and

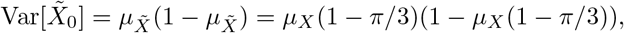

which concludes the proof.

#### 5.2 Proof of Lemma 2.6

Let *Z*_*S*_ = (*X*^(1)^(*S*), *X*^(2)^(*S*)) be a S-IBN process based on the process of standing up times *S*, with parameters *µ, σ, α*_1_, *α*_2_, *β*_1_, *β*_2_, *π*_1_, *π*_2_. Let *Y*^(1)^ := *Y*^(1)^(*S*_1_), *Y*^(2)^ := *Y*^(2)^(*S*_2_) be two independent IBNs with the same parameters, i.e., *µ, σ, α*_1_, *β*_1_, *π*_1_, and *µ, σ, α*_2_, *β*_2_, *π*_2_ respectively, based on two independent processes of standing up, *S*_1_ and *S*_2_. For *B* ∼ Ber(*u*) and independent from all other random variables, let

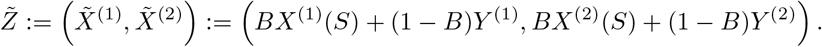

Thus,

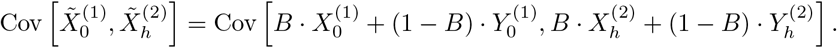

We multiply and get for the individual terms, with 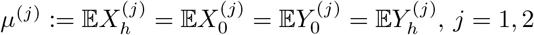

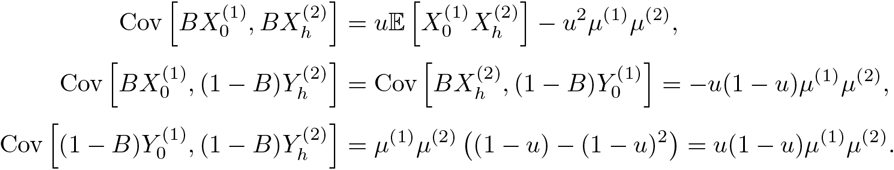

We thus have

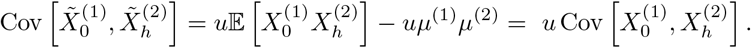

With Var[*X*^(*i*)^] = Var[*Y*^(*i*)^] for *i* = 1, 2 the assertion follows for 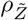.

#### 5.3 Details on the parameter estimation in the IBN and S-IBN

As described in Section 2.5, the distribution of observed SCs in the IBN and S-IBN are mixture distributions of (1) non-interrupted SCs and of (2) the first and (3) second part of interrupted SCs. As we do not know which observed SC comes from which distribution, and because successive observed SCs are no longer independent, maximum likelihood estimation would be extremely cumbersome.

Therefore, we propose to fit the theoretical PDFs of SCs, *f*_*SC*_, and LFs, *f*_*LF*_, to their empirically observed histograms. The theoretical PDFs of the mixture distributions of SCs and LFs are derived below in Lemma 5.1 and 5.2.

In our fitting algorithm, we used *κ* = 20 equally sized bins for the histogram of standing cycle (SC) lengths for each animal *j*. Additional simulation studies showed no notable improvement in model fit or parameter estimation with higher bin counts, indicating that 20 bins provide an effective balance between resolution and computational efficiency. Thus, the bin sizes for the SCs for each animal *j* had a bin size of 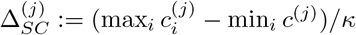, where 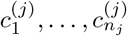 denote the lengths of all fully observed SCs in animal *j*. For the histogram of LFs, we proceeded analogously, i.e., 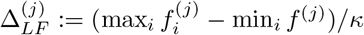 where 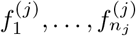 denote the LFs of all fully observed SCs in animal *j*. This yielded bin-midpoints 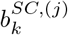 and 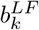, *k* = 1, …, *κ*, as

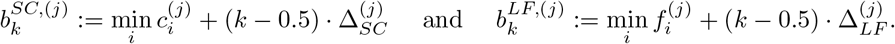

The corresponding bin counts, normed to a unit area of the histogram, are then 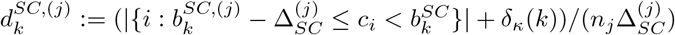, where “*<*“ is replaced by “≤” for *k* = *κ*, and analogously for 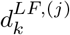.

Then *f*_*SC*_ and *f*_*LF*_ are fitted jointly using a non-linear-least squares approach, i.e. parameters are estimated by minimizing the term

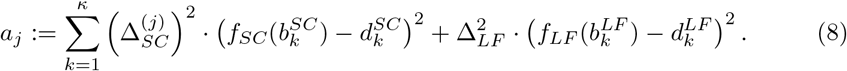

The parameters in the S-IBN are estimated analogously, i.e., we first estimate the parameters *µ, σ, α*_1_, *α*_2_, *β*_1_, *β*_2_, *π*_1_, and *π*_2_ by minimizing the term 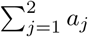. In order to estimate the parameters *δ* and *u*, we predict the CCF 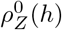 as derived in Theorem 2.5 with the estimated parameters for an S-IBN with *u* = 1, *δ* = 0 and then fit the function *u · ρ*_*Z*_(*h* + *δ*) to the observed CCF using nonlinear least squares.

Now we derive the theoretical densities *f*_*SC*_ and *f*_*LF*_. Intuitively, both the distributions of SCs and of LFs are necessarily mixture distributions of observations from non-interrupted standing cycles and of interrupted standing cycles. If each cycle is interrupted with probability *π* and then produces two cycles, the fraction of uninterrupted cycles is (1 − *π*)*/*(1 + *π*).

##### Lemma 5.1 (PDF of Standing Cycles in the IBN)

*The probability density function f*_*SC*_ *of the standing cycles in an IBN*(*µ, σ*^2^, *α, β, π*) *is given by*

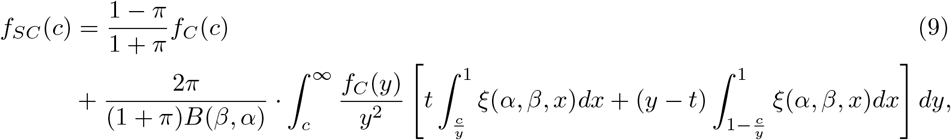

*where f*_*C*_ *is the PDF of 𝒩* _+_(*µ, σ*), *B is the beta function and ξ*(*α, β, x*) := *x*^*β*−3^(1 − *x*)^*α*−1^.

To derive the PDF *f*_*SC*_ of the observed SC length in the IBN model with interruption probability *π*, we note that *f*_*SC*_ can be written as the finite mixture

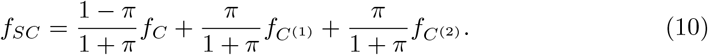

In case of an interruption, the two standing cycles 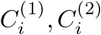 can be written as (see Figure 12)

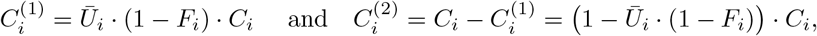

with *Ū*_*i*_ := max{*U*_*i*1_, *U*_*i*2_}. Hence

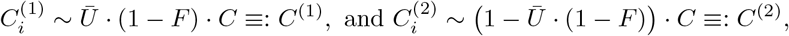

with *F* ∼ Beta(*α, β*), *C* ∼ *𝒩* _+_(*µ, σ*) for all *i* ∈ ℤ.

**Fig 12.**
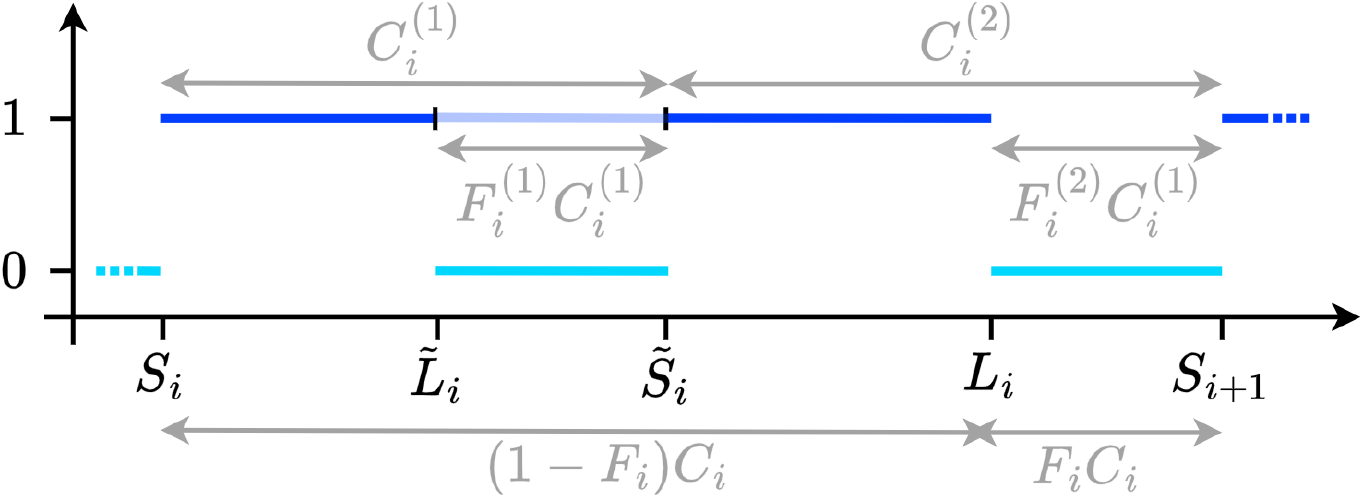
Illustration of notation and variables used in the proofs of Lemmas 5.1 and 5.2.

Since for a random variable *F* on [0, 1] with PDF *f*_*F*_ it holds that *f*_1−*F*_ (*t*) = *f*_*F*_ (1 − *t*) it follows that

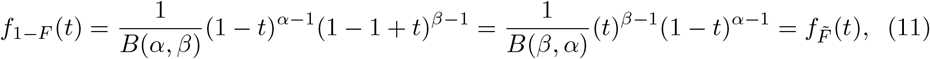

with 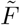 ∼ Beta(*β, α*) and *B*(*α, β*) = *B*(*β, α*) the beta function.

In order to derive the density of *Ū ·* (1 − *F*), we use first that for two independent random variables *X, Y* with PDFs *f*_*X*_, *f*_*Y*_, the PDF of the product *XY* is given by

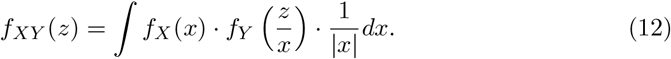

Further, we use that the PDFs *f*_*Ū*_ and *f*_*U*_ of the maximum *Ū*:= max{*U*_1_, *U*_2_} and minimum *<u>U</u>* := min{*U*_1_, *U*_2_} of two independent uniform random variables *U*_1_, *U*_2_ ∼ Unif[0, 1] are

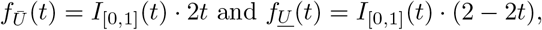

with expected values 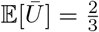 and 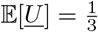.

Applying (12) and (11) yields

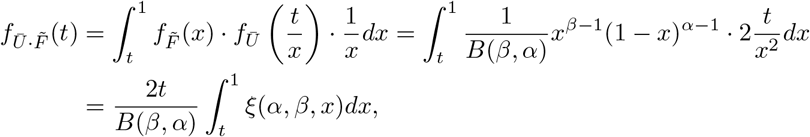

with

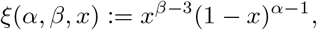

for *t* ∈ [0, 1] and 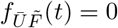 for *t* ∉ [0, 1]. Applying (12) again yields

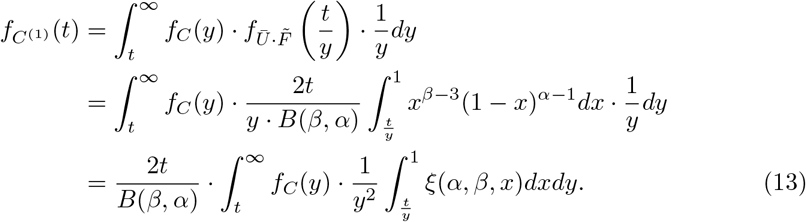

To derive the density of *C*^(2)^, note that *Ū ·* 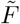 takes values in [0, 1]. Therefore the PDF 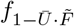 of 1 − *Ū ·* 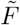 obeys 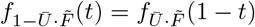, for *t* ∈ [0, 1] and 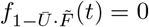 for *t* ∉ [0, 1]. Thus, it follows that

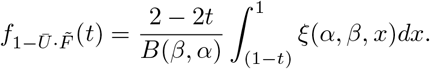

This leads to

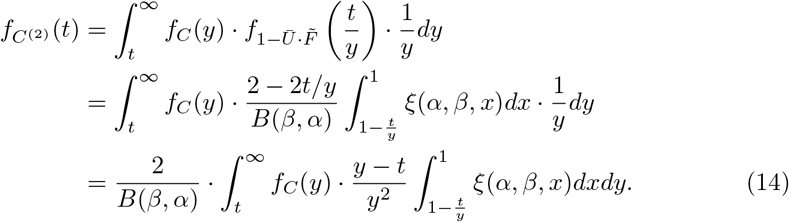

Plugging in (13) and (14) into (10) yields equation (9).

##### Lemma 5.2 (PDF of Lying Fractions in the IBN)

*The probability density function f*_*LF*_ *of the lying fractions in an IBN*(*µ, σ*^2^, *α, β, π*) *is given by*

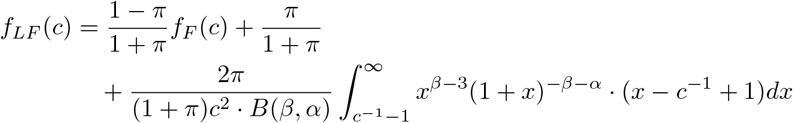

*for c* ∈ [0, 1] *and f*_*LF*_ (*c*) = 0 *for c* ∉ [0, 1], *where f*_*F*_ *is the PDF of Beta*(*α, β*) *and B is the beta function*.

We will follow a similar outline as in Lemma 5.1. The PDF *f*_*LF*_ of the observed lying fractions in the IBN with interruption probability *π* can be written as the finite mixture

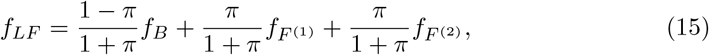

with *f*_*F*_ as the PDF of Beta(*α, β*). In case of an interruption, the two lying fractions 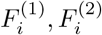 can be written as (see Figure 12)

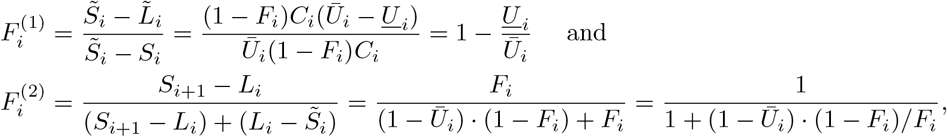

with *<u>U</u>*_*i*_ := min{*U*_*i*1_, *U*_*i*2_} and *Ū*_*i*_ := max{*U*_*i*1_, *U*_*i*2_}. Hence, with *<u>U</u>* and *Ū* defined analogously and as in Appendix 5.3,

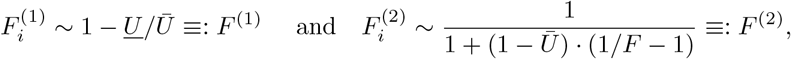

with *F* ∼ Beta(*α, β*) for all *i* ∈ ℤ.

For the density of *F*^(1)^ we note that given *Ū, <u>U</u>* is uniformly distributed on [0, *Ū*]. Therefore, both *<u>U</u>/Ū* and *F*^(1)^ = 1 − *<u>U</u>/Ū* are uniformly distributed on the unit interval, such that

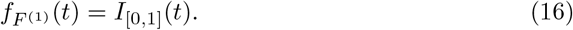

For the density of *F*^(2)^ we note that if *F* ∼ Beta(*α, β*) then 1*/F* − 1 ∼ Beta′(*β, α*) where Beta′ denotes the Beta prime distribution with density

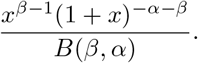

Using the fact that 1 − *Ū* ∼ *<u>U</u>* we get

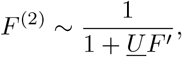

where *F*′ ∼ Beta′(*β, α*). The density of *UF*′ is calculated as

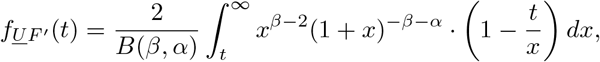

and the density of 1 + *UF*′ is given by *f*_1+*<u>U</u>·F*′_ (*t*) = *f*_*<u>U</u>·F*′_ (*t* − 1). Finally, we note that for non-negative random variables *X >* 0, it holds that 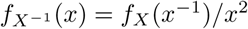 for all *t* ∈ [0, 1]. Thus, we get

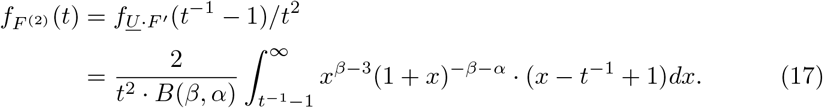

Plugging in (16) and (17) into (15) concludes the proof.

## Supporting information

**S1 Fig**.

---

**S1. Fig.**
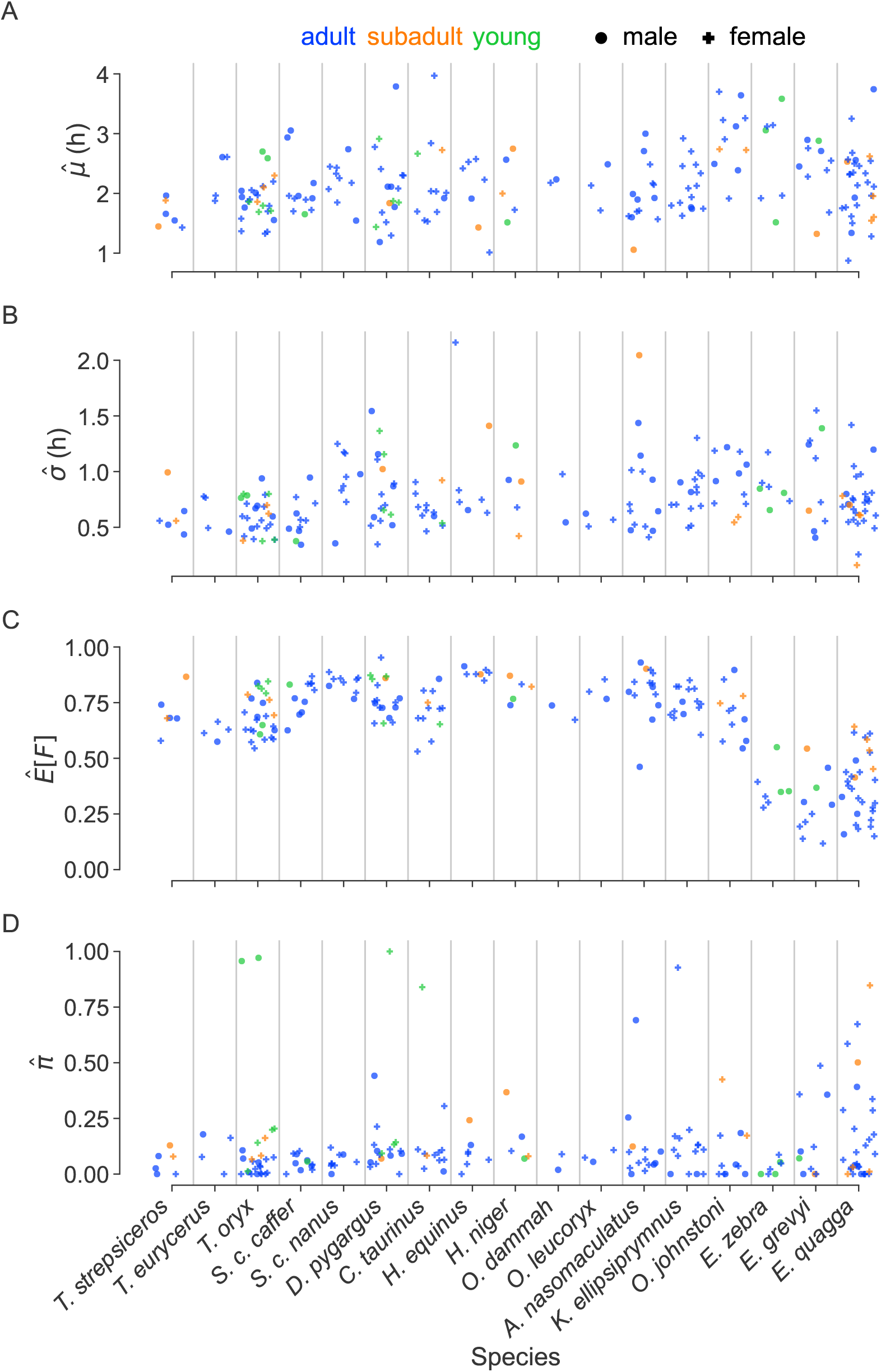
Parameter estimates in the IBN for all 198 animals.

## References

1. IUCN Red List of Threatened Species; 2022. Available from: https://www.iucnredlist.org.

2. Rose P, Robert R. Evaluating the activity patterns and enclosure usage of a little-studied zoo species, the sitatunga (Tragelaphus spekii). Journal of Zoo and Aquarium Research. 2013;1(1):14–19. doi:10.19227/jzar.v1i1.12.

3. Berger A. Activity patterns, chronobiology and the assessment of stress and welfare in zoo and wild animals. International Zoo Yearbook. 2010;45(1):80–90. doi:10.1111/j.1748-1090.2010.00121.x.

4. Brando S, Buchanan-Smith HM. The 24/7 approach to promoting optimal welfare for captive wild animals. Behavioural processes. 2018;156:83–95. doi:10.1016/j.beproc.2017.09.010.

5. Walsh B, Binding S, Holmes L. While you were sleeping Zooquaria. 2019;105:28–29.

6. Rose PE, Riley LM. Conducting Behavioural Research in the Zoo: A Guide to Ten Important Methods, Concepts and Theories. Journal of Zoological and Botanical Gardens. 2021;2(3):421–444. doi:10.3390/jzbg2030031.

7. Caboń-Raczyńska K, Krasińska M, Krasiński Z. Behaviour and daily activity rhythm of European bison in winter. Acta Theriologica. 1983;28:273–299. doi:10.4098/at.arch.83-22.

8. Packard JM, Loonam KE, Arkenberg CR, Boostrom HM, Cloutier TL, Enriquez EJ, et al. Behavioural profiles of African bovids (Hippotraginae). Journal of Zoo and Aquarium Research. 2014;2(3):8287. doi:10.19227/jzar.v2i3.74.

9. Reta R, Solomon Y. Diurnal activity patterns of Burchells zebra (Equus quagga, Gray 1824) in Yabello Wildlife Sanctuary, Southern Ethiopia. International Journal of Biodiversity and Conservation. 2014;6(8):616–623. doi:10.5897/ijbc2014.0736.

10. Zhang E. Daytime activity budgets of the Chinese water deer. Mammalia. 2000;64(2):163–172. doi:10.1515/mamm.2000.64.2.163.

11. Leuthold B, Leuthold W. Daytime activity patterns of gerenuk and giraffe in Tsavo National Park, Kenya. African Journal of Ecology. 1978;16(4):231–243. doi:10.1111/j.1365-2028.1978.tb00444.x.

12. Bennie JJ, Duffy JP, Inger R, Gaston KJ. Biogeography of time partitioning in mammals. Proceedings of the National Academy of Sciences of the United States of America. 2014;111(38):13727–13732. doi:10.1073/pnas.1216063110.

13. Gravett N, Bhagwandin A, Sutcliffe R, Landen K, Chase MJ, Lyamin OI, et al. Inactivity/sleep in two wild free-roaming African elephant matriarchs - Does large body size make elephants the shortest mammalian sleepers? PLoS ONE. 2017;12(3):e0171903. doi:10.1371/journal.pone.0171903.

14. Wu Y, Wang H, Wang H, Feng J. Arms race of temporal partitioning between carnivorous and herbivorous mammals. Scientific Reports. 2018;8(1):1713. doi:10.1038/s41598-018-20098-6.

15. Burger AL, Fennessy J, Fennessy S, Dierkes PW. Nightly selection of resting sites and group behavior reveal antipredator strategies in giraffe. Ecology and Evolution. 2020;10(6):2917–2927. doi:10.1002/ece3.6106.

16. Sicks F. REM sleep as indicator for stress in giraffes (Giraffa camelopardalis). Mammalian Biology. 2016;81:16. doi:10.1016/j.mambio.2016.07.052.

17. Tobler I, Schwierin B. Behavioural sleep in the giraffe (Giraffa camelopardalis) in a zoological garden. Journal of Sleep Research. 1996;5(1):21–32. doi:10.1046/j.1365-2869.1996.00010.x.

18. Hahn-Klimroth M, Kapetanopoulos T, Gübert J, Dierkes PW. Deep learning-based pose estimation for African ungulates in zoos. Ecology and Evolution. 2021;11(11):6015–6032. doi:10.1002/ece3.7367.

19. Gübert J, Hahn-Klimroth M, Dierkes PW. BOVIDS: A deep learning-based software package for pose estimation to evaluate nightly behavior and its application to common elands (Tragelaphus oryx) in zoos. Ecology and Evolution. 2022;12(3):e8701. doi:10.1002/ece3.8701.

20. Gübert J, Hahn-Klimroth M, Dierkes PW. A large-scale study on the nocturnal behavior of African ungulates in zoos and its influencing factors. Frontiers in Ethology. 2023;2:1219977. doi:10.3389/fetho.2023.1219977.

21. Gübert J, Schneider G, Hahn-Klimroth M, Dierkes PW. Nocturnal behavioral patterns of African ungulates in zoos. Ecology and Evolution. 2023;DOI: 10.1002/ece3.10777.

22. Ryder OA, Feistner ATC. Research in zoos: A growth area in conservation. Biodiversity and Conservation. 1995;4(6):671–677. doi:10.1007/BF00222522.

23. Dement WC, Kleitman N. Cyclic variations in EEG during sleep and their relation to eye movements, bodily motility and dreaming. Electroencephalography and Clinical Neurophysiology. 1957;9:673–690.

24. Gübert J, Hahn-Klimroth M, Dierkes PW. BOVIDS: A deep learning-based software package for pose estimation to evaluate nightly behavior and its application to common elands (Tragelaphus oryx) in zoos. Ecology and Evolution. 2022;12(3).

25. Møller J. Shot noise Cox processes. Adv Appl prob. 2003;35(3):614–640.

26. Møller J, Torrisi GL. Generalised shot noise Cox processes. Adv Appl prob. 2005;37(1):48–74.

27. Schneider G. Messages of oscillatory correlograms - a spike train model. Neural Comput. 2008;20(5):1211–1238.

